# Histone Tail Electrostatics Modulate E2-E3 Enzyme Dynamics: A Gateway to Regulate Ubiquitination Machinery

**DOI:** 10.1101/2022.08.27.505537

**Authors:** Dineli T. S. Ranathunga, Hedieh Torabifard

## Abstract

BRCA1 (BReast Cancer-Associated protein 1), a human tumor suppressor, plays a key role in genome stability and DNA repair. Heterodimerization of BRCA1 with BARD1 is important for its stability, maximal Ub ligase (E3) activity and cooperative activation of UbcH5c (E2). Recent studies demonstrate the importance of ubiquitination of the nucleosomal H2A C-terminal tail by BRCA1/BARD1-UbcH5c (E3-E2) in which its mutations inhibit ubiquitination, predispose cells to chromosomal instability and greatly increase the likelihood of breast and ovarian cancer development. Due to the lack of molecular-level insight on the flexible and disordered H2A C-tail, its ubiquitination mechanism by BRCA1/BARD1-UbcH5c and its function and relationship to cancer susceptibility remain elusive. Here, we use molecular dynamics simulations to provide molecular-level insights into the dynamics of the less-studied H2A C-tail and BRCA1/BARD1-UbcH5c on the nucleosome surface. Our results precisely identify the key interactions and residues that trigger conformational transitions of BRCA1/BARD1-UbcH5c, and characterize the important role of histone electrostatics in their dynamics. We show that the dynamics of the H2A C-tail, combined with the highly mobile UbcH5c, define the ubiquitination capacity. Furthermore, our data demonstrate a mechanistic basis for the probability of ubiquitination of C-tail lysines in the ordered and disordered regions. Altogether, the findings of this study will provide unrevealed insights into the mechanism of H2A C-tail ubiquitination and help us understand the communication between E2-E3 enzymes and nucleosome to regulate ubiquitination machinery, paving the way for the development of effective treatments for cancer and chronic pain.

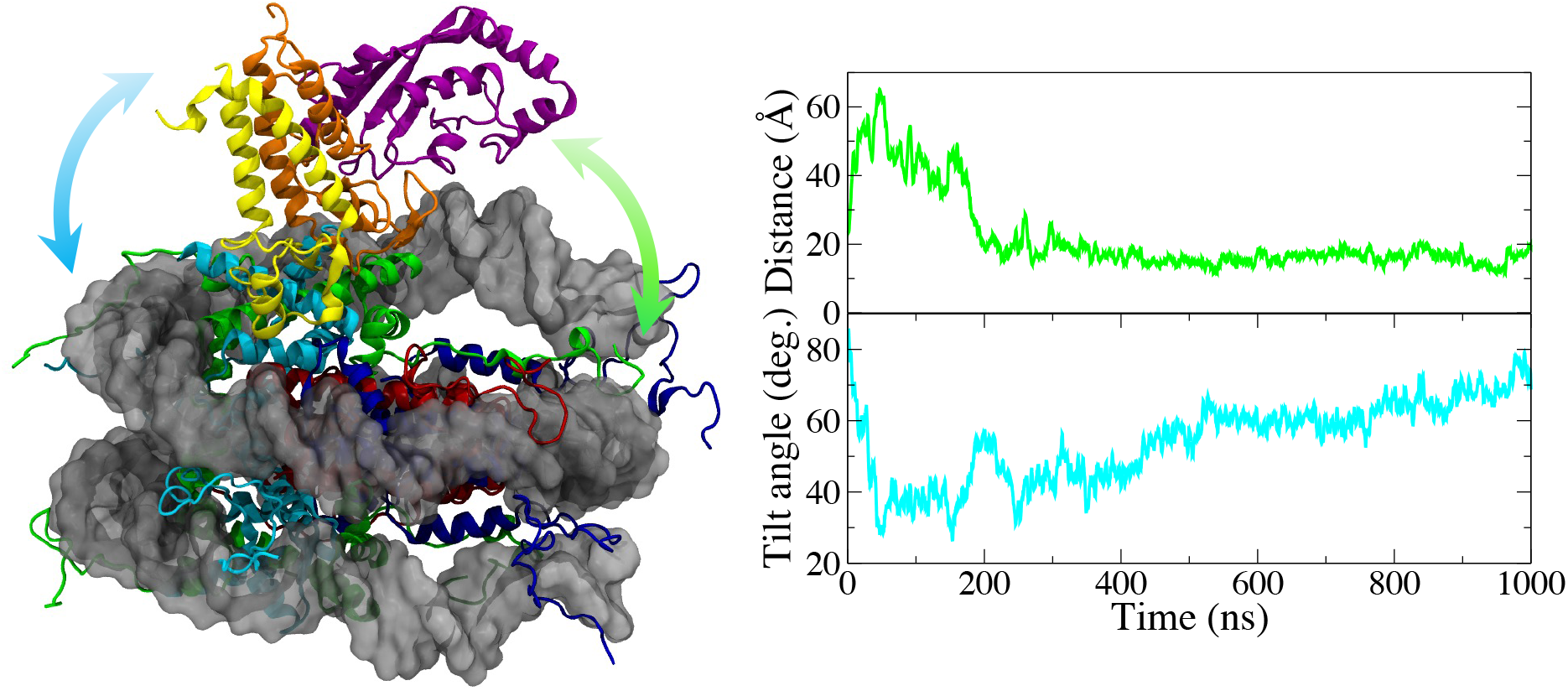

## INTRODUCTION

Nucleosome core particles (NCPs) are the fundamental repeating units of chromatin packaged within the cell nucleus. Every human cell contains millions of NCPs. ^1–4^ A single NCP consists of 147 base pairs (bps) of DNA sequence wrapped around a protein core. ^4–9^ The protein core forms an octameric structure with two copies of four histone proteins (H2A, H2B, H3 and H4). ^4^ The structure of a NCP is shown in Figure 1. Each of the histone proteins possesses characteristic N-terminal and C-terminal tails, which are densely packed with basic residues such as lysine and arginine and are capable of interacting more directly with DNA and various proteins within the nucleus.^6,10^ These histone tails are susceptible to extensive covalent post-translational modifications (PTMs), such as ubiquitination, methylation, acetylation, and phosphorylation, which will affect their structure, dynamics and function. These modifications can alter DNA-histone contacts and their conformation, and consequently affect their intrinsic properties, chromatin organization, and underlying transcriptional processes.^6,10–15^ The landscape of known histone PTMs that influence the structural and functional state of chromosomes, and the chromatin-based processes such as gene expression, transcription, DNA replication, and DNA repair, continues to expand.

**Figure 1:**
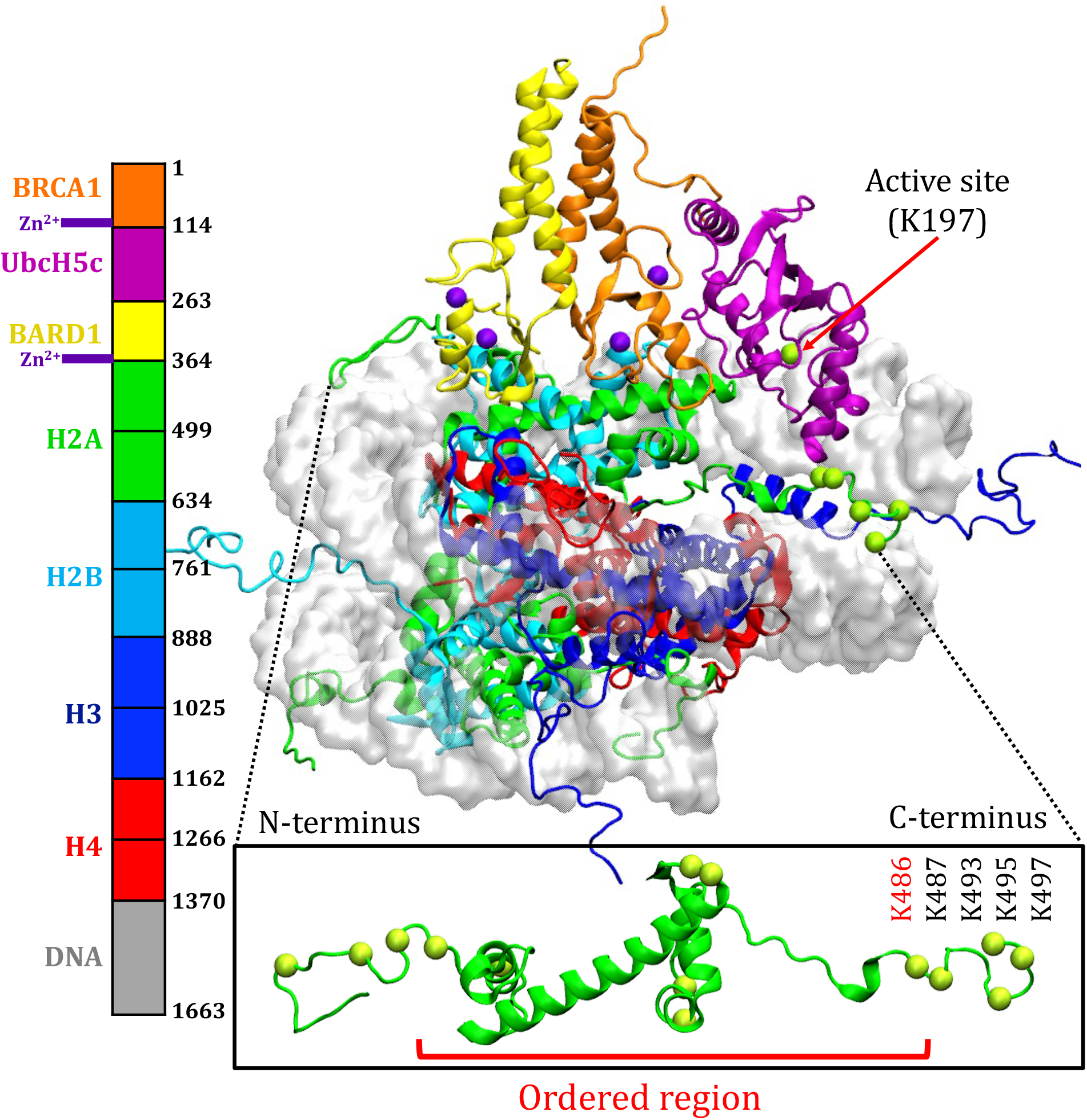
BRCA1/BARD1-UbcH5c/NCP simulated structure and full H2A chain with lysine C_*α*_ atoms shown in green spheres. Lysine residues in the ordered C-tail of H2A are labeled in red and lysine residues in the disordered C-tail of H2A are labeled in black.

In cancer, misregulation of chromatin-based processes leads to inappropriate inactivation of tumor suppressors. ^16^ Indeed, PTMs at histone tails are central to the biology of several cancers. For instance, ubiquitination of the H2A histone tails is a prominent histone PTM that has been shown to be altered in various types of cancer, including breast, ovarian, and lung cancer^6,12,17,18^, and thus has received intense attention in recent literature.^6,12,17,19–23^ Ubiquitination is a process of binding ubiquitin (Ub), that regulates many essential cellular processes. All ubiquitination pathways share three common steps shown in Figure S1 involving 3 enzymes (E1, E2 and E3). BRCA1 (BReast Cancer-Associated protein 1) expressed in breast cells and other tissues is a human tumor suppressor that is primarily responsible for repairing damaged DNA and is well known for its E3 Ub ligase activity.^6,12,22,24^ The BRCA1 and its obligatory partner BARD1 (BRCA1-associated RING domain protein 1) form a large heterodimeric complex with a 4 helix bundle dimerization interface which contains 4 tetrahedrally-coordinated zinc metal centers. This BRCA1/BARD1 hetero-complex possesses enhanced E3 activity, with a significant role in repairing the damaged DNA.^12,19,25^ Mutations in BRCA1/BARD1 have been shown to increase the risk of familial breast and ovarian cancer and are associated with various sporadic cancers. ^12,19,26,27^ Recently, a few targets for E3 Ub ligase activity of BRCA1/BARD1 have been identified *in vivo* and nucleosomal histone H2A was discovered as one of the bona fide substrates for BRCA1/BARD1-dependent E3 ligase activity. ^28^ BRCA1/BARD1 together with UbcH5c specifically ubiquitinates lysine residues in the H2A C-terminal tail^17^. UbcH5c belongs to the family of ubiquitin-conjugating enzymes and is one of two mammalian E2 proteins that display BRCA1-dependent ligase activity. ^29,30^ It covalently binds to BRCA1.^31^ Brzovic *et al*.^29^ reports that a mutation in the UbcH5c-BRCA1 interface is sufficient to severely reduce or eliminate BRCA1/BARD1 E3 activity. Therefore, the integrity of BRCA1/BARD1-UbcH5c is critical for its ability to transfer Ub to the amino (-NH_2_) group of lysine residues on the H2A C-terminal tail. ^12,19,32,33^

Very recently, the cryo-EM structures of the BRCA1/BARD1-UbcH5c/NCP complex have been characterized.^12,19^ Although the disordered histone tails in these cryo-EM structures are not characterized due to their extremely dynamic nature, it is proposed that ubiquitination occurs at lysine residues residing in the fully disordered C-terminal tail of H2A. ^12,19^ Figure 1 shows the structure of H2A and the location of the disordered H2A C-tail. Previous studies revealed that BRCA1/BARD1 RING heterodimers are topologically similar and share several functional characteristics with other E3 ubiquitin ligase enzymes such as Ring1b/Bmi1 and RNF168.^12,17,19^ Therefore, the BRCA1/BARD1 ubiquitination mechanism of H2A is proposed compared to these E3 enzymes.^12,19^ In the cryo-EM structure of BRCA1/BARD1-UbcH5c, the distance between the active site of UbcH5c and K486 in the ordered region of H2A (Figure 1) is larger (19 Å) compared to the distance in Ring1b/Bmi1 system (9 Å).^19^ Therefore, Witus *et al*.^19^ proposed that this BRCA1/BARD1-UbcH5c arrangement can lift UbcH5c from the histone surface to distance its active site from the H2A lysine residues in the ordered region. This position disfavors the transfer of Ub to lysines in the ordered region and promotes the modification of last three lysines in the disordered H2A C-tail, *i*.*e*. K493, K495 and K497. ^19^ Recent *in vitro* and *in vivo* mutation studies have revealed that BRCA1/BARD1-UbcH5c can modify one or more of these three lysine residues.^12,17,19^ This is in contrast to the ubiquitination of H2A by other E3 ligase complexes such as Ring1b/Bmi1 which mainly targets K486/K487 in the H2A C-tail. ^5,22^ Indeed, the majority of H2A C-tail ubiquitination reported so far occurs at K487 by different E3 ligase enzymes. ^34^

Currently, very few data are available on the mechanism of ubiquitination of the H2A C-tail lysine substrate by BRCA1/BARD1-UbcH5c and some of the results on the selectivity of H2A ubiquitination remain controversial. For instance, a study reported that K493 ubiquitinates with the highest efficiency^19^, while another study shows that only K495 and K497 ubiquitinate^17^. Additionally, the intrinsically disordered histone tails can adopt a broad range of conformations upon binding to other proteins or DNA. Hence, the characterization of their conformational dynamics and interactions with DNA and PTM enzymes are extremely challenging.^12,19,35–41^ Furthermore, the lack of molecular-level mechanistic information on BRCA1/BARD1-UbcH5c ubiquitination of H2A hinders our understanding of the role of BRCA1/BARD1-UbcH5c and its relationship to cancer. Therefore, it is critical to explore molecular-level mechanism of H2A C-tail ubiquitination to elucidate the underlying basis of the experimental observations. All-atom molecular dynamics (AA-MD) simulation is a promising approach to elucidate molecular-level details of the conformational ensemble of proteins with intrinsically disordered regions and can be used to provide insight into molecular mechanisms.^24,42,43^ Moreover, to precisely understand how histone H2A C-tail dynamics regulate the ubiquitination machinery, it is necessary to model the complete structure of histone in the nucleosome architecture. Therefore, in this work, we modeled the full-length structure of histones in the NCP using AA-MD simulations. We investigated the global conformational dynamics of BRCA1/BARD1 and UbcH5c on the NCP surface, and identified the residues and the interactions that trigger these movements. To gain insight into the impact of histone tail dynamics on long-range communication between PTM elements (*i*.*e*. DNA, PTM enzymes, histones), electrostatic interactions were studied by calculating the interaction energies of mutants and wild type (WT). Moreover, the selectivity and the efficiency of ubiquitination of H2A C-tail lysine by BRCA1/BARD1-UbcH5c were characterized. Overall, this study provides a qualitative picture of the mechanism of ubiquitination of the H2A C-tail by BRCA1/BARD1-conjugated UbcH5c on the NCP surface in a fully human system. The findings of this research will allow us to understand the key electrostatic interactions to control communication between E3-E2 and different PTM elements, and to design potential therapeutics.

## METHODOLOGY AND SIMULATION DETAILS

### Model preparation

To build the BRCA1/BARD1-UbcH5c/NCP complex, a cryo-EM structure of the BRCA1/BARD1UbcH5c/NCP was obtained from the protein data bank (PDB ID 7JZV^19^ with resolution 3.9 Å). The 7JZV nucleosome model does not contain histone tails and only contains 139 bp of DNA. The swiss model^44^ web server was used to search for homology modeling templates for all eight histone tails, and the target sequences were threaded onto the templates. The structure of the missing histone tail segments and the missing DNA bps were constructed using the 1KX5^45^ PDB model (resolution 1.9 Å) of the nucleosome core particles as a reference. High sequence identity indicates the high accuracy of the model. The sequence identities between a modeled protein and the templates are shown in Table S1. The final structure contains BRCA1 bound to Ubiquitin-conjugating enzyme UbcH5c (E2), BARD1 and a NCP (residue IDs are shown in Figure 1). Protonation states of the titratable residues were assigned using PropKa3.0^46^ and H++^47^. The protein chains were terminated with ACE (acetylated) and NME (amidated) patches. The systems were solvated with TIP3P water^48^ and explicitly neutralized using 150 mM KCl that extended at least 10 Å from the solute^49^, creating systems of ∼400,000 atoms (Figure S2). The overall solvation box of 149.37 × 169.28 × 194.35 Å^3^ (Figure S2) was made periodic in all three directions.

Subsequently, various mutant models were generated using the AMBER Leap program^50^ to study the electrostatic effects of H2A C-tail (mutating all H2A C-tail lysine residues to alanine (all-ALA) and arginine (all-ARG)) and the ubiquitination selectivity (single mutants of K486A, K487A, K493A, K495A and K497A). Sixteen total AA-MD simulations were carried out on WT and all mutants (including multiple trial runs). In the current study, the active site of UbcH5c was modeled as a lysine (K197) consistent with the cryo-EM^19^ and the NMR^5^ experiments instead of a cysteine. According to Witus *et al*.^19^, this cysteine to lysine mutation of the UbcH5c active site increases the association of the BRCA1/BARD1UbcH5c/NCP complex and stabilizes the interactions. Finally, all the systems were subjected to AA-MD simulations.

### Simulation details

AA-MD simulations of the WT and mutated BRCA1/BARD1-UbcH5c/NCP systems, including the full-length structures of the histones (with tails) were performed. Five independent WT simulation runs were conducted, starting from the same initial conformation with different initial velocities to confirm the reproducibility of results and improve the sampling. Similarly, three independent trajectories were simulated for all alanine (all-ALA) and all arginine (all-ARG) mutated systems. Only one MD simulation was conducted for each single mutant system (K486A, K487A, K493A, K495A and K497A). All the models were first energy minimized to remove steric clashes. For all structures first 2000 steepest descent minimization steps were performed before switching to 3000 steps of conjugate gradient minimization. During the initial minimization the protein and nucleosome heavy atoms were harmonically restrained by a force constant of 500 kcal*/*mol*/*Å^2^. This first step was followed by an energy minimization for 10,000 steps without any constraints. After minimization, the systems were gradually heated up to 300 K using Langevin dynamics with a collision frequency of 1.0 ps^−1^ in the NVT ensemble for 1.5 ns constraining the backbone coordinates with a harmonic potential (k=10 kcal*/*mol*/*Å^2^). Then, the systems were equilibrated at 300 K for 2 ns under constant pressure conditions (NPT, 1 atm). Once the systems achieved the targeted temperature, production simulations were performed at 1 atm pressure and 300 K temperature using Langevin dynamics with a collision frequency of 1.0 ps^−1^ under NPT ensemble with the Langevin thermostat and Berendsen barostat. The production length of each simulation is given in Table S2, and snapshots were saved every 20 ps. Initially, all of the MD runs were conducted for 500 ns, but some selected trials were extended to a maximum of 1100 ns to assess the convergence of simulations and to sample all the major conformational dynamics of BRCA1/BARD1-UbcH5c on the NCP surface, minimizing computational cost. The cumulative simulation time in this study is 10.55 *µ*s. The simulations were performed with a timestep of 2 fs and a 12 Å cutoff for non-bonded interactions. During the calculations, all bonds involving hydrogen atoms were constrained with the SHAKE algorithm ^51^ and the particle mesh Ewald (PME) method^52^ was employed to treat long-range Coulomb interactions.

All the simulations were performed using the GPU-accelerated version of the pmemd program in the AMBER20^50^ software package with Amber ff14sb^53^ and DNA.OL15^54^ parameters to describe the proteins and the nucleic acids, respectively. Zinc AMBER force field (ZAFF) developed by Peters *et al*.^55^ was used to represent tetrahedrally-coordinated zinc metal centers that are bound to the nearby cysteine and histidine residues. The trajectories were visualized with the VMD program.^30^ All the analyses including root mean square deviation (RMSD) and fluctuation (RMSF), hydrogen bond, linear interaction energy (LIE), tilt angle, principal component analysis (PCA), and distance analysis on trajectories were carried out using the CPPTRAJ module^56^ available in the AMBER20 suite. Histograms were generated using CPPTRAJ where the population sum was normalized to 1.0 to present the results of all MD trials (for WT and mutants) as normalized probability distributions. Contact frequency calculations were performed using an in-house TCL code ^49,57^ and energy decomposition analysis (EDA) was used to calculate non-bonded inter-molecular interaction energies (Coulomb and vdW interactions) between selected residues using a FORTRAN90 program^58–60^.

## RESULTS AND DISCUSSION

### Conformational dynamics and stability

First, five independent simulations were performed on WT to increase sampling, and all five trajectories were analyzed together as described in the simulation details section to both understand conformational stability and identify conformational dynamics. The structural stability convergence of each MD simulation was examined using RMSD of the protein domains and DNA, to ensure the adequacy of the simulation time and the validity of the analysis. Figure S3 shows the RMSDs of the domains in BRCA1/BARD1-UbcH5c/NCP relative to the initial configuration. RMSD of the BRCA1/BARD1-UbcH5c entire domain, DNA and nucleosome core containing 8 histones with (Histone core w/ tails) and without (Histone core w/o tails) tails were first calculated in Figures S3a and S3b. The overall RMSD results illustrate that the simulations are well equilibrated (Figure S3a). Since the disordered regions can have heterogeneous ensembles of conformations, the nucleosome core with histone tails displays significantly higher degree of conformational changes than the nucleosome core when histone tails are not considered (only considering the ordered regions) (Figures S3a and S3b).

The RMSD around 6 Å of the entire BRCA1/BARD1-UbcH5c domain indicates a large structural change during the simulations (Figure S3a). To further explore this, the RMSDs of individual domains of UbcH5c, BRCA1 and BARD1 for WT trajectories were calculated and results are presented in Figure 2a and Figures S3(c-e). Compared to BRCA1 and BARD1, UbcH5c shows relatively smaller RMSDs (Figure 2a). The calculation of RMSF of individual residues in UbcH5c is shown in Figure 2b. Highlighted regions in the UbcH5c structure (shown in mint green) show regions with RMSF *>*5 Å. These regions include the area containing the UbcH5c active site and an unstructured loop area containing residues 228-260, which have large structural contributions to the movement of the entire UbcH5c domain. In contrast to UbcH5c, the RMSDs of BRCA1 and BARD1 show broad populations, indicating that both BRCA1 and BARD1 conformations deviate more from their original structures during the simulations (Figures 2a and S3). The RMSF data in Figures 2c and 2d highlight the unstructured protein loops and terminal regions of BARD1 and BRCA1 (shown in mint green), where relatively high structural fluctuations are discovered. Altogether, the results suggest that BRCA1/BARD1-UbcH5c displays large conformational dynamics on the NCP surface.

**Figure 2:**
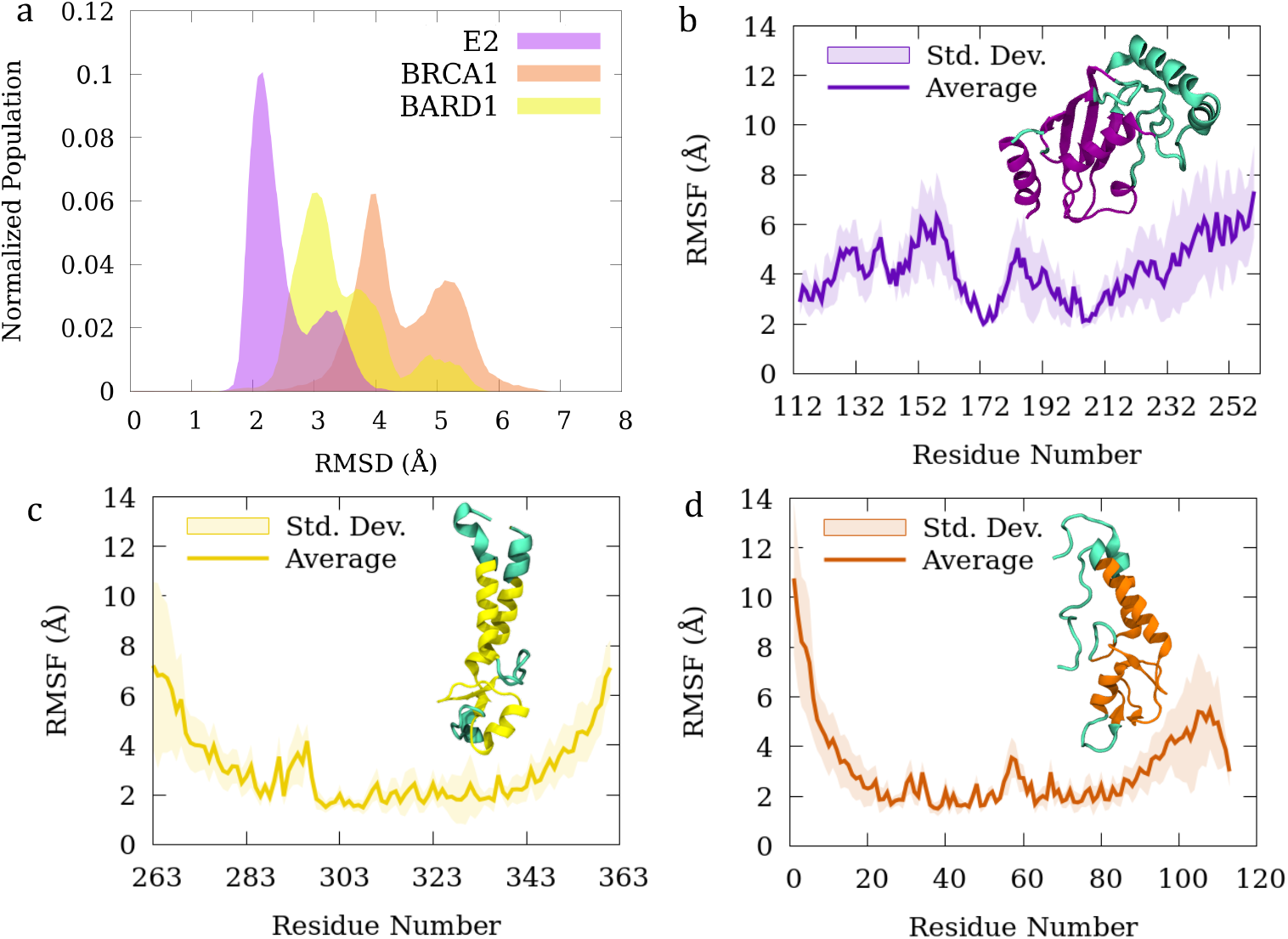
a. Normalized backbone root mean square deviation population of UbcH5c (E2), BRCA1, and BARD1 domains of all WT trajectories with respect to the initial conformation. Individual trial results are shown in Figure S3c-e. Side chain root mean square fluctuations of b. UbcH5c, c. BARD and d. BRCA1. The regions with higher (*>*3.5 Å) fluctuation are colored in mint green in BARD1 and BRCA1. For UbcH5c the regions with *>*5 Å RMSF are colored in mint green.

### PCA reveals the entire BRCA1/BARD1-UbcH5c conformational space

PCA of protein configurations of MD trajectories were performed to quantify the underlying dynamics of BRCA1/BARD1-UbcH5c on the NCP surface. The results of the WT trajectories are shown in Figures 3 and S4-S6. The proportion of variance versus principal components (PCs) presented in Figure S4a shows the contribution of each PC in the overall dynamics. On average, for the WT simulations, the first six PCs represent *>*80% of the overall conformational variability of the entire BRCA1/BARD1-UbcH5c domain in the MD simulations (Figures S4a). Skjaerven *et al*.^61^ reports that if there is sufficient sampling, the nature of the individual PCs calculated in one MD run will be reproduced by another MD run where the only difference between the MD runs is the set of initial velocities, however, there will not always be a one-to-one correspondence between a given mode and its corresponding PC in different MD runs.^61^ In agreement with Skjaerven *et al*.^61^ and confirming sufficient sampling, we noted that the principal components between different MD seeds overlap considerably and only the order of the modes were permuted due to randomness and different simulation times (Figure S5). This suggests that in the current study, all WT trajectories describe similar types of motions. To describe the structural fluctuations in BRCA1/BARD1 and UbcH5c that occurred during the MD simulations, the first six principal components calculated for the WT simulations were explored and six different modes (A-F) corresponding to PCs were shown in Video S1.

**Figure 3:**
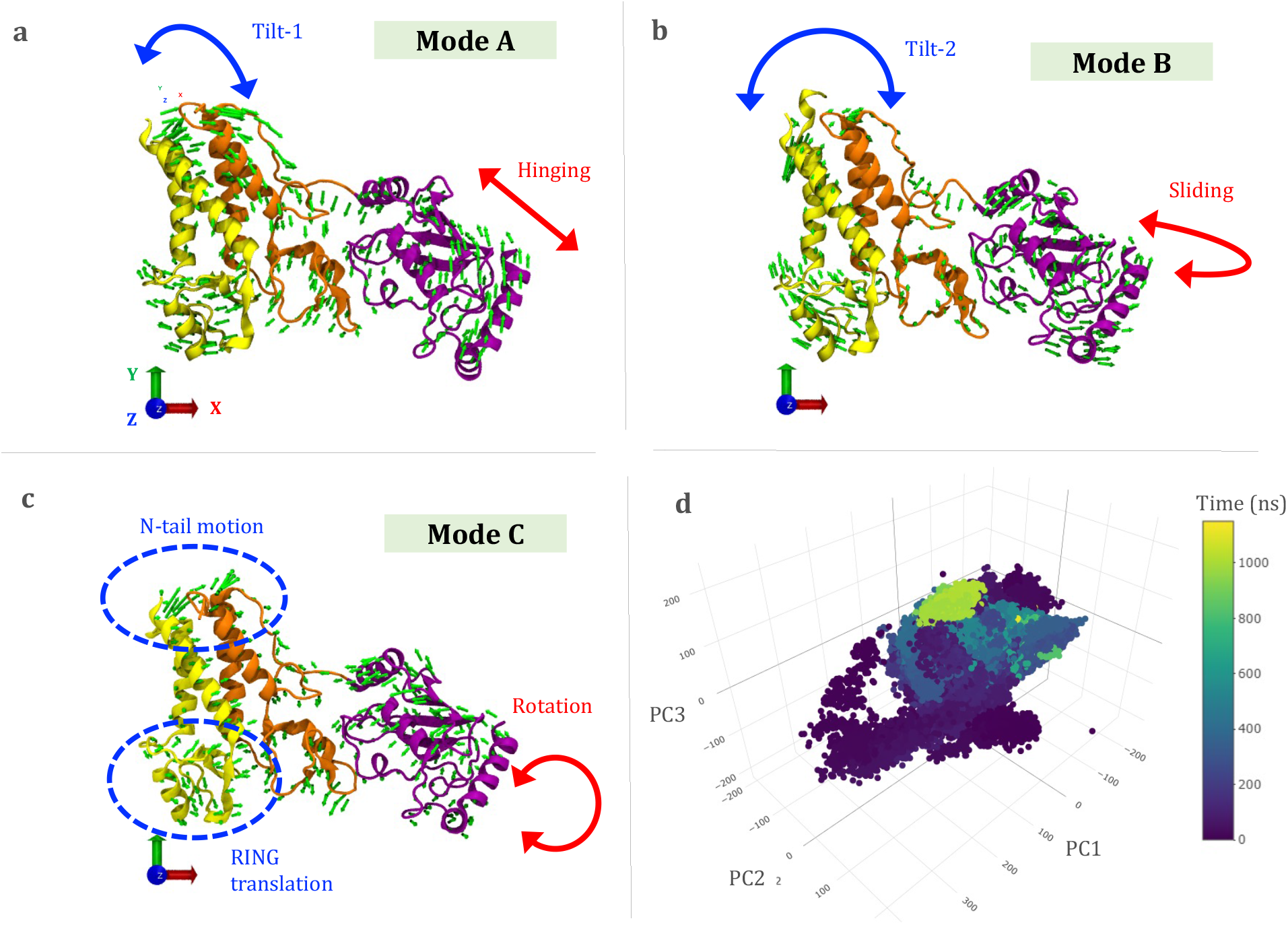
a,b,c. Vector field representation of the principal components (PCs) obtained from WT simulations for the entire BRCA1/BARD1-UbcH5c domain. For ease of visualization, only BRCA1/BARD1-UbcH5c are shown here. The NCP is positioned parallel to the XZ plane on the trajectories. d. Conformational space sampled in WT simulations as a function of first three principal components (PC1, PC2, PC3) obtained from the analyses of five WT trajectories colored according to time to differentiate the clustering with time and structural stability convergence. Molecules are colored as follows: BRCA1 (orange), BARD1 (yellow) and UbcH5c (purple).

Each mode containing multiple elements and the same element repeating in multiple modes were detected when comparing the vector field representations of the first 6 PCs (shown in Video S1 and Figure S5). In Figure 3 only the first 3 modes that show the highest probability of occurrence (probabilities are given in Figure S5) are presented, illustrating all the large amplitude motion elements. The first mode (A) shows a UbcH5c hinging motion perpendicular to the NCP surface (hinging motion) and BRCA1/BARD1 tilting toward H2B (Tilt-1 motion) (Figure 3a and Video S1). The second mode (B) represents the sliding of UbcH5c (sliding motion) on the NCP surface parallel to the surface and the Tilt-2 movement of BRCA1/BARD1 (parallel to XY plane) (Figures 3b and Video S1). The third component (mode C) exhibits the sliding and hinging movements of UbcH5c at a smaller amplitude, along with some rotation of UbcH5c around its principal axis (Figure 3c). This movement is defined as the rotation motion in the current study. Mode C also shows more tilt of BRCA1/BARD1 in the H2B direction, with large translational fluctuations of the RING domain of BARD1 in the XY plane direction (called RING translation motion) and movement at the N-terminus of BARD1 (called N-tail motion). These large N-terminal tail fluctuations also occur in mode B and are noted in the RMSF data in Figure 2. These motions (N-tail motion and RING translation motion) have a large impact on BRCA1/BARD1 tilt. The fourth principal component mode (mode D) contains multiple moving elements, including high-amplitude UbcH5c sliding motion along with BRCA1/BARD1 leaning toward H2B, and the translation of BARD1 RING domain (Figure S5 and Video S1). Mode E primarily shows movement at the N-terminus of BARD1, which skews BRCA1/BARD1. Mode F represents the hinging movement of UbcH5c and the Tilt-1 movement of BRCA1/BARD1 at a large amplitude (Figure S5) similar to mode A. However, the direction of the movements of BRCA1/BARD1-UbcH5c is different in mode F compared to mode A (Video S1). Mode F shows a seesaw-like movement of BRCA1/BARD1-UbcH5c that is consistent with the behavior proposed by Witus *et al*.^19^

Generally, in PCA, the transitions displayed by a single PC are not exactly how the system experiences motion; indeed the actual motion of the system during the simulation is always represented by the combination of PCs.^50,56,62^ Therefore, in the current study, we focused on all of the above large-amplitude movements of the BRCA1/BARD1 and UbcH5c domains to accurately describe the conformational transitions. All identified global movements of BRCA1/BARD1 and UbcH5c domains, *i*.*e* Tilt-1, Tilt-2, hinging and sliding movements compared to their initial orientations, are shown in Figure S6. To provide a more quantitative comparison of PCs, the structures of backbone atoms in all WT MD trajectories are projected onto essential space (planes) defined by PC1/PC2/PC3 in Figure 3d since all observed global backbone changes in the ensemble are well described by the first 3 PCs. This allows visualization of the sampled conformational space during MD calculations. In Figure 3d, each point represents one conformation saved during the MD simulations, and the density of points represents the population of conformations sampled in the MD trajectories. The magnitude of motion along PC2 (PC2 ranges from -200 to 200) and PC3 (PC3 ranges from -200 to 150) is relatively smaller than that along PC1 (PC1 ranges from -350 to 350) (Figures 3d and S4b). Figure 3d shows that the largest motion and majority of structural variations occur within the first 200 ns of the WT simulations. In addition, all the trajectories converge at the end (after 200 ns) and the conformations are populated mostly around (0,0,0) (Figures 3d and S4b). Taken together, BRCA1/BARD1 and UbcH5c dominant motions on the NCP surface starting from any given configuration were identified as BRCA1/BARD1 Tilt-1, Tilt2, N-tail motion and RING translation motions and UbcH5c hinging, sliding and rotation motions (Figures 3 and S6).

### R73 and W319 retain BRCA1/BARD1 on the NCP surface

To further investigate the aforementioned flexible states accessible to BRCA1/BARD1- UbcH5c within the surface of the nucleosome, residues at the interface that are responsible for holding BRCA1/BARD1-UbcH5c on the surface were first characterized. Contact frequency analysis data in Figure S7 shows that the BRCA1 and BARD1 RING domain regions include residues 51, 54-57, 69-77, 301-304, 316-334, and 336 -338 (highlighted in mint green in Figure S7) remain within 7 Å to the NCP surface for more than 95% of the simulation time. Experimental mutagenesis studies^12,19^ reported that when residues R73 in the BRCA1 RING domain and W329 in the BARD1 RING domain are mutated to alanine, BRCA1/BARD1 binding to the nucleosome surface was undetectable and also showed an attenuating effect of H2A ubiquitination. Consistent with experiments, our results suggest that R73 and W329 can act as anchoring motifs to keep BRCA1/BARD1 on the surface, which can also serve as pivot points around which BRCA1/BARD1 rotates. In 20% of the WT simulations, the loop containing W329 moved along H2B and formed an adjacent interface with residues in the *α*3- and *α*C-helices of H2B (helices are shown Figure S8a) due to the BARD1 RING translation motion (Figure 3c), while R73 remained trapped on the NCP surface for 100% of the simulation time. The hydrogen bond calculation results in Figure S9 identify the residues in the NCP binding pockets that interact with R73 and W329 and remain within 7 Å of these two residues throughout the simulations. Among these residues, the H2A and H2B residues bind strongly to the BRCA1 and BARD1 RING domains, helping to retain BRCA1 and BARD1 on the surface. Overall, the results demonstrate that these nucleosome-binding factors confine BRCA1 and BARD1 to the NCP surface and may allow characteristic movements of the BRCA1/BARD1-UbcH5c complex on the surface.

### BRCA1/BARD1 tilting mechanism

Of all the flexible states accessible to BRCA1/BARD1 within the nucleosome surface around the above discussed nucleosome-binding factors, Tilt-1 motion appears at the highest amplitude (Mode A in Figure S5). Therefore, to provide a quantitative view of the Tilt-1 motion, the angle between one of the principal axes (z) passing through the center of mass of one of the BARD1 (or BRCA1) *α*-helices and the vector passing through the H2B *α*C-helix was calculated using the dot product and the normalized Tilt-1 angle population data of all WT trajectories are presented in Figures 4a and S10a. The initial angles of BARD1 and BRCA1 relative to H2B in the cryo-EM structure are about 84 and 86 degrees, respectively. Angular populations propagating up to 20 degrees during the simulation confirm the tilting movement of BRCA1/BARD1 towards H2B (Figures 4 and S10). Although these tilt angles initially decrease, they increase over time and BRCA1/BARD1 returns to its initial position (Figures 4b and S10b). This return motion is further corroborated by the time variation of the tilt angle of WT MD run 1 shown in Figures S11a-b.

**Figure 4:**
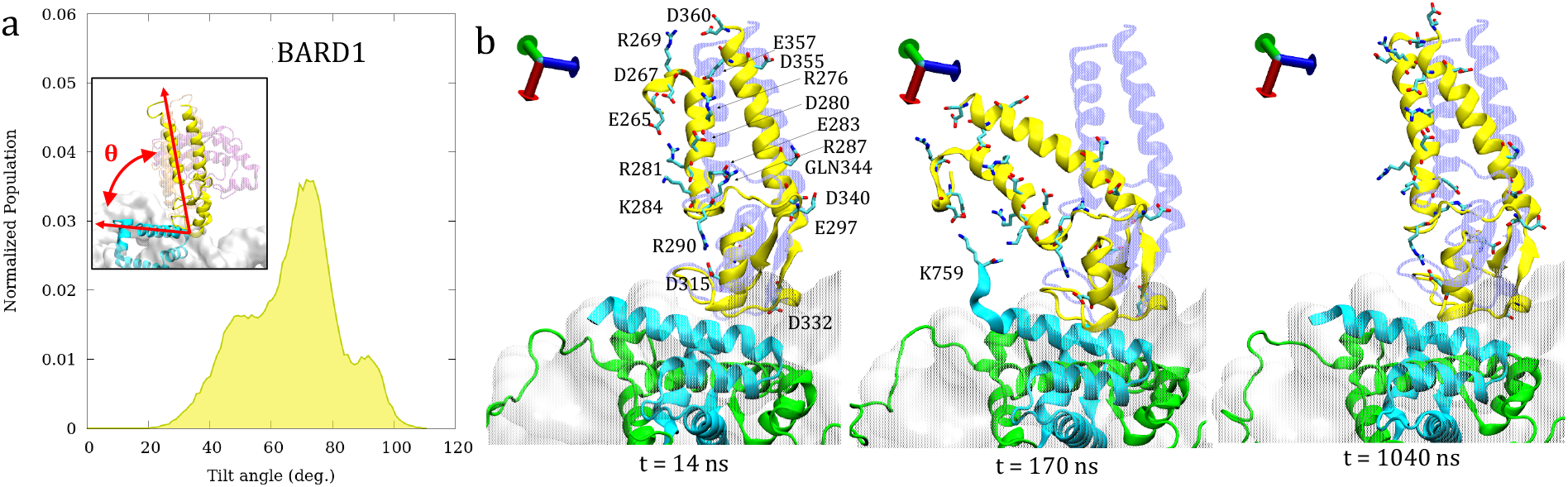
a. BARD1 tilt angle distribution for WT trajectories. b. The directional Tilt-1 movement of BARD1 relative to its initial configuration. Residues in BARD1 contribute to the tilting motion (that show electrostatic energy higher than -100 kcal/mol with H2B and H2A in Figures S12a and S12c) are labeled. BRCA1 results are shown in Figure S10. Coloring as follows: H2B (cyan), H2A (green), UbcH5c (transparent purple), BARD1 (yellow), DNA (gray), initial structure (transparent blue), carbon atoms (cyan), oxygen (red), and nitrogen (dark blue).

Figures 4b and S10b show the directional tilt movements of BARD1 and BRCA1 relative to the initial configuration (shown in transparent blue), respectively. To gain a qualitative understanding of the tilting mechanism, the effect of residues in the domains of interest was investigated. The EDA results in Figure S12 show BARD1 and BRCA1 residues that strongly interact with H2B and H2A. The MD trajectories reveal that initially E265 and D267 residues at the upper N-terminus of BARD1 show interactions with positively charged residues in the helical region of BARD1 (Figures 4b and S13). These attractive interactions between the charged residues out-compete the stabilization of the initial BARD1 *α*-helical structure near the N-terminus (residues in region 264-271 shown in the trajectory snapshot at t=14 ns in Figure 4b) and result in N-tail motion (Figure 3). Subsequently, BARD1 N-terminal residues, along with charged residues (mostly negatively charged residues) in the helical region (Figures 4b at 170 ns and S12), strongly interact with the oppositely charged residues in the H2B (such as K759) and H2A to advance the tilt. Due to these strong electrostatic interactions, BARD1 remained tilted for a period of time during the simulations (Figure S11a). However, over time BARD1 leans back to its original position due to the forces from the UbcH5c side. This return movement of BARD1 causes the N-terminus residues to fold back into their helical structure (Figure 4b at 1040 ns).

In addition to the dynamic N-terminal movement, PCA reveals a translational movement of the RING domain of BARD1 (Figure 3, RING translational motion) that also greatly aids its Tilt-1 movement. The main residues involved in this motion are located in the unstructured loop region 322-333 (the enlarged region in Figure S7), which shows a large standard deviation of RMSF in Figure 2c, due to its flexible nature. Simulation trajectories reveal that when this low-amplitude translational movement of the BARD1 RING domain occurs, it affects the movement of BRCA1 on the NCP surface, pushing basic residues in the BRCA1 RING domain toward the DNA. These interactions are responsible for the attractive energy of both negatively charged BRCA1 and DNA shown in Figure 5c (shown in light red). The net negative charges of BRCA1 and DNA are shown in Figure S8a. However, this translational mobility of BRCA1 RING domain on the NCP surface is markedly lower than that of BARD1 due to the strong interaction of the R73 anchor with the NCP acid patch (Figure S9). Nonetheless, the tilting mechanism (Tilt-1 and Til-2) of BRCA1 is equivalent to that of BARD1. Similar to BARD1, the conformational dynamics of the disordered N-tail of BRCA1 influences the tilting motion. The most interacting residues of BRCA1 with H2B and H2A are shown in Figures S10b, S12b and S12d. Taken together, the results suggest a strong contribution of the charged residues on BRCA1/BARD1 tilting motions.

**Figure 5:**
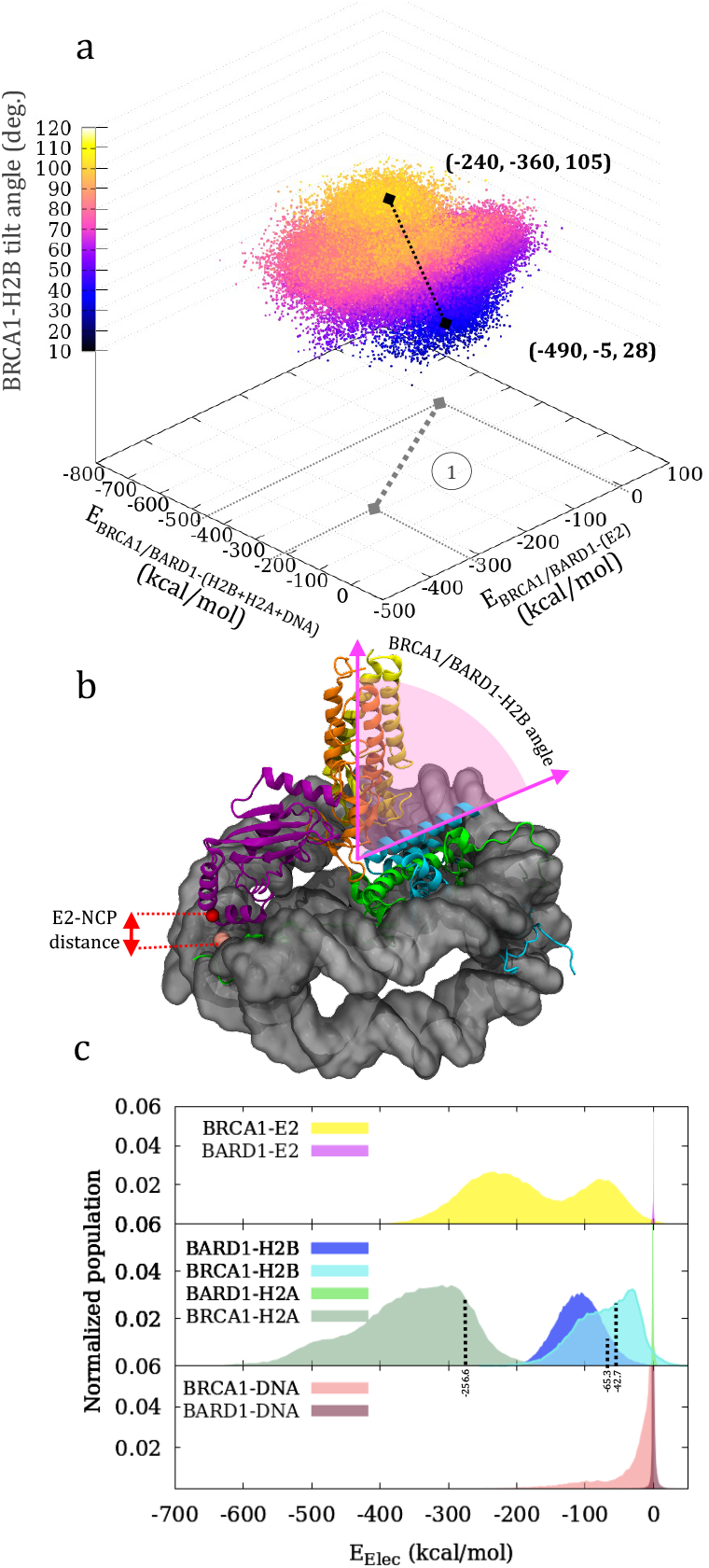
a. Results of the total non-bonded interaction energy between BRCA1/BARD1 with H2B, H2A and DNA (X axis), the total non-bonded interaction energy between BRCA1/BARD1 with UbcH5c (Y axis), the BRCA1-H2B tilt angle (colored Z axis). To explain the correlations between the data, two points were selected from the clustered data points and their projection on a plane is marked. The coordinates of the points follow the (E_BRCA1*/*BARD1−(H2B*/*H2A*/*DNA)_, E_BRCA1*/*BARD1−(E2)_, BRCA1 − H2B angle) order. b. Atoms selected to measure the BRCA1-H2B tilt angle and the E2-NCP distance parameters in the current study. The BRCA1-H2B tilt angle was measured using the dot product of vectors passing through the H2B *α*C-helix and principal component axis passing through BRCA1. The distance between E242 (C_*α*_) on UbcH5c (E2) and 1443 (P) on DNA was considered as the UbcH5c (E2)-NCP distance. Coloring as follows: H2B (cyan), H2A (green), UbcH5c (purple), BARD1 (yellow), BRCA1 (orange), DNA (dark gray). c. BRCA1 and BARD1 electrostatic interaction energy with E2, H2B, H2A, and DNA for WT.

### H2B and H2A electrostatic interactions drive BRCA1/BARD1 tilting motions

In order to characterize the driving force for the tilting behavior, the interaction energies of BRCA1/BARD1 with the NCP domains were explored and their effect on the BRCA1-H2B angle was studied. Figure 5a, show the correlation between 3 parameters, the total nonbonded interaction energy between BARD1/BRCA1 with H2B, H2A and DNA (x-axis), the total non-bonded interaction energy between BARD1/BRCA1 and UbcH5c (E2) (y-axis), and tilt angle between BRCA1 and H2B (colored z-axis). Figure 5b shows the selected vector angle for BRCA1-H2B tilt angle calculations. To explain the correlations between the parameters, two points (from the clustered data) were selected (Figure 5a). The data demonstrate that when tilt angle decreases (from 105° to 28°), BRCA1/BARD1 interactions with H2B+H2A+DNA increase (from -240 kcal/mol to -490 kcal/mol) and interaction energy between BRCA1/BARD1 and UbcH5c decreases (from -360 kcal/mol to -5 kcal/mol). This suggests that the interactions from both sides can be involved in the determination of BRCA1/BARD1 movement (Figure 5a correlation 1).

To identify which component has the largest contribution to BRCA1/BARD1 movement, we investigated the individual interaction energy between BRCA1/BARD1 and other components. Since the electrostatic contribution to the total non-bonded interaction energy is much higher than the van der Waals contribution (Figure S14a), we only showed the electrostatic interaction energies of BRCA1 and BARD1 with UbcH5c (E2), H2B, H2A and DNA in Figure 5c. The results show that BRCA1/BARD1 interactions with UbcH5c are mainly due to the electrostatic interactions between BRCA1 and UbcH5c (Figure 5c). This is a result of the covalent attachment of BRCA1 to UbcH5c that keeps the RING domain of BRCA1 always close to UbcH5c. However, these UbcH5c interactions show a strong effect on BRCA1/BARD1 return movement after tilting toward the NCP surface. Therefore, in Figure 5a, as the tilt angle increases, BRCA1/BARD1 approaches UbcH5c, the energy of BRCA1/BARD1 with UbcH5c increases. The net charges of H2B (+17.99e) and H2A (+16.99e) are fairly similar (Figure S8a), and both H2A and H2B residues were found to interact with BARD1 and BRCA1 (Figure S12). Figure 5c shows that BRCA1 and BARD1 strongly interact with H2B and H2A compared to DNA. These electrostatic interactions are mainly caused by the BRCA1 and BARD1 RING domains. In addition to adhering to the NCP surface, some regions of the RING domains show tilting behavior (Figures 4b and S10b). Therefore, these strong electrostatic interactions of BRCA1/BARD1 with H2B/H2A correspond to holding BRCA1/BARD1 on the NCP surface and their tilting movement. The energy changes compared to the initial interaction energies (marked with dashed lines) shown in Figure 5c further confirm the contribution of H2B and H2A to the movement of BRAC1 and BARD1. Moreover, Figure S14b shows that the BARD1-H2B interaction energy changes upto -100.5 kcal/mol and the BRCA1-H2B energy changes upto -99.6 kcal/mol when the tilt angle varies between 40° and 20° indicating the involvement of the BRCA1/BARD1 helical region in the tilting motion. This also reveals that helical domain interactions are dominant towards H2B. Altogether, these findings elucidate that electrostatic interactions from the H2A and H2B primarily drive the tilt of the BARD1/BRCA1 heterocomplex toward NCP surface, and UbcH5c interactions show a strong effect on BRCA1/BARD1 return movement. This forward and backward tilting motion and interactions of BRCA1/BARD1 may confer the conformational space accessible for UbcH5c on the nucleosome surface.

### H2A C-tail affects the movement of UbcH5c on the NCP surface

In section 3.2, PCA showed that UbcH5c can undergo two large-scale movements at the NCP surface: 1. hinging movement perpendicular to the nucleosome surface and 2. sliding motion parallel to the surface of the nucleosome (Figure S6). To quantify these movements, the relative distance between the residue D242 (C_*α*_) on UbcH5c and a selected stable DNA location (P atom of residue 1443) was calculated (Figure 5b), and these E2-NCP distances are reported in Figure 6a. The initial E2-NCP distance in the cryo-EM structure is 12.83 Å. The change in distance from 2 Å to 68 Å confirms the movement of UbcH5c with a higher probability of moving about 42 Å from the NCP surface (Figure 6a). Considering all WT simulation runs, this articulation behavior of UbcH5c was also identified as an event independent of time. The broadly distributed C_*α*_ − C_*α*_ distances between the active site of UbcH5c and H2A C-tail lysine residues shown in Figure 6b further confirm this conformational transition of UbcH5c connected to BRCA1/BARD1 on the surface of the nucleosome which is consistent with recently proposed mechanism by Witus *et al*.^19^

**Figure 6:**
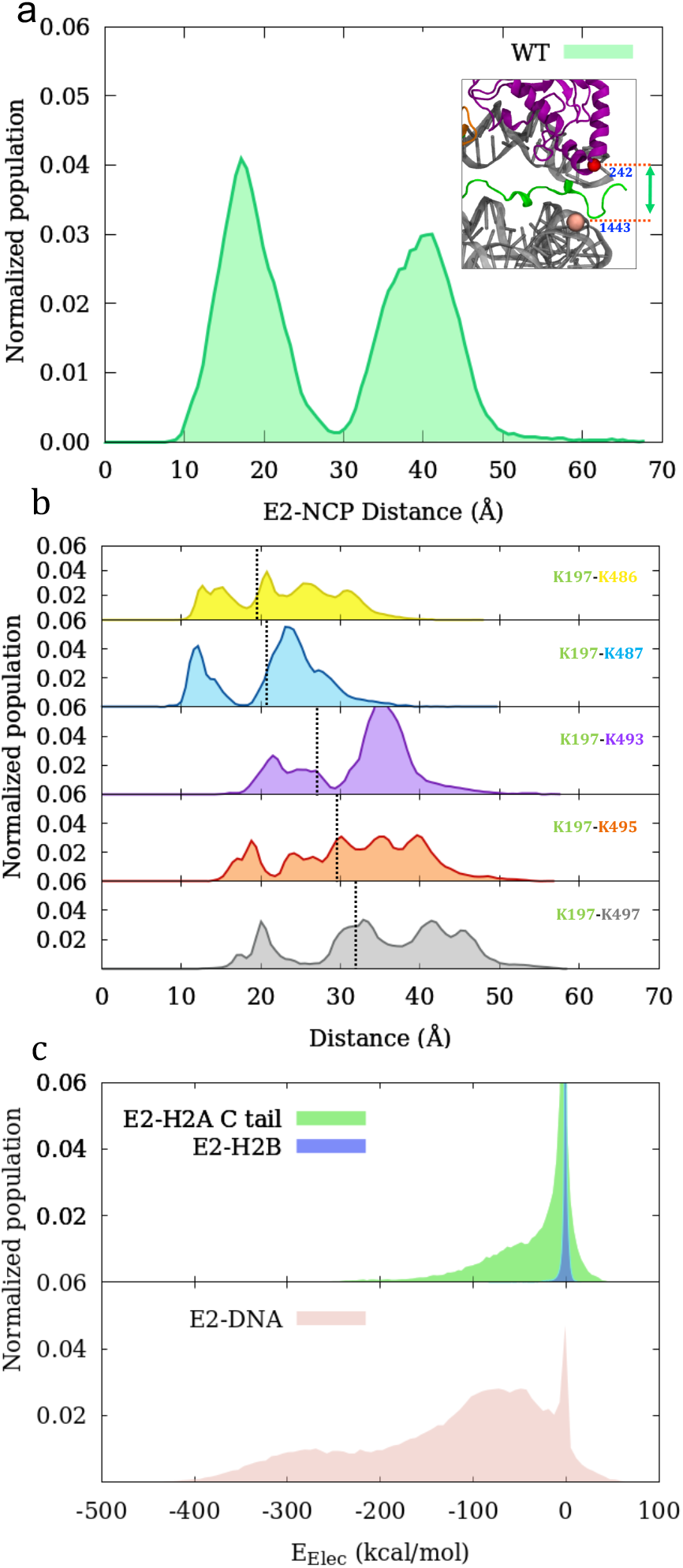
a. UbcH5c (E2)-NCP distance for all WT trajectories. E2-NCP distance is calculated using the residue D242 (C_*α*_) on UbcH5c and 1443 (P atom) on DNA. b. C_*α*_-C_*α*_ distance between UbcH5c active site (K197) and H2A C-tail lysine residues. Initial distances in the cryo-EM structure are shown in dotted lines. c. UbcH5c (E2) electrostatic interaction energy with H2A C-terminal tail, H2B and DNA for WT.

To understand the main causes of the UbcH5c movement on the NCP surface, we focused on the interactions acting on UbcH5c. The electrostatic interaction energies of UbcH5c with surrounding domains presented in the Figures 5c and 6c show that the H2A C-tail, DNA, and BRCA1 have strong electrostatic interactions with UbcH5c that may contribute to UbcH5c motion. The interactions between E2 and H2A N-tail and ordered region are negligible compared to that with H2A C-tail. Once UbcH5c moves away from the surface, its orientation (due to its rotational motion in Figure 3c) is such that some charged residues face the NCP surface and strongly interact with oppositely charged residues on the H2A Ctail and DNA (causing strong electrostatic attraction energy in Figure 6c). Contact frequency analysis identifies the residues in BRCA1/UbcH5c that are most frequently found within 7 Å of the 5 lysine residues in the H2A C-tail and DNA over the entire WT simulation time and the results are highlighted in Figures 7 and S8b. Among the residues in the highlighted regions, acidic residues (such as D228, D229, E234, D242, D244, and E252) are oriented in a direction where they can interact especially with the disordered H2A C-terminal lysine residues (basic) (Figure 7) and some basic residue were found in close proximity to the DNA (Figure S8b). These residues play a critical role in the return or downward movement of UbcH5c.

**Figure 7:**
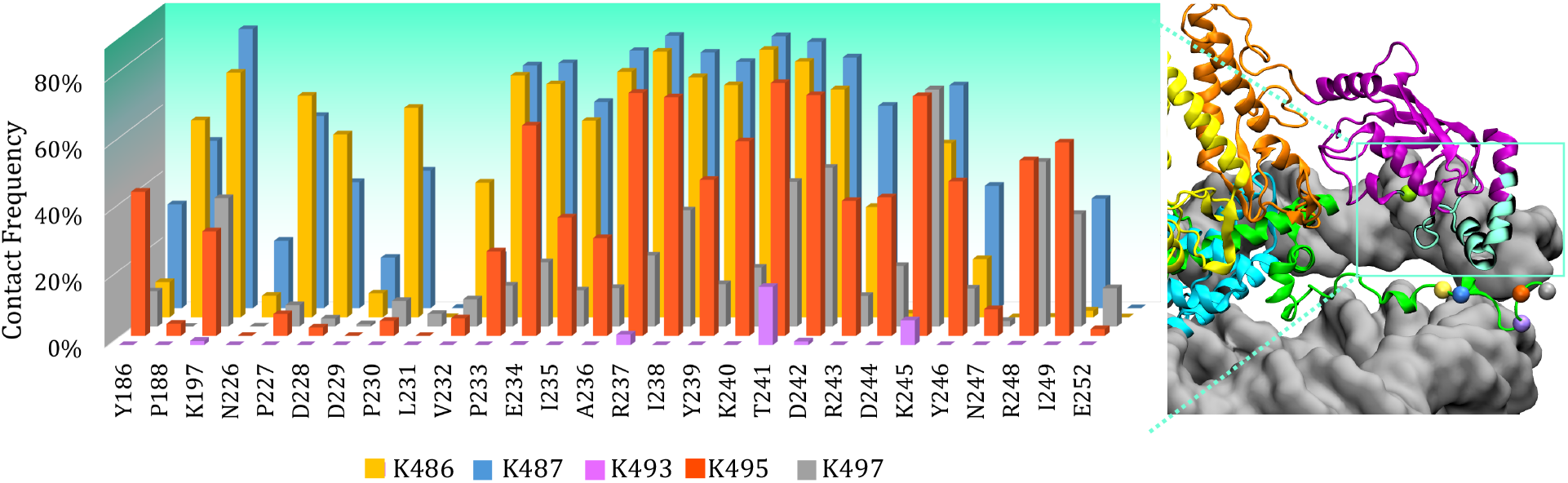
Contact frequency for WT simulations. Percentage of time that residues in BRCA1/UbcH5c can be found close (*<*7 Å) to lysine residues in H2A C-tail. Coloring as follows: DNA (dark gray), BARD1 (yellow), BRCA1 (orange), UbcH5c (purple), H2B (cyan), H2A (green) and UbcH5c region that contacts H2A C-tail/DNA is colored in mint green. C_*α*_ atoms of H2A C-tail lysine residues are labeled with the same colors in the left panel. Contact frequency results of DNA are shown in Figure S8b.

In addition to H2A and DNA, BRCA1 also shows strong electrostatic attraction energy to UbcH5c in Figure 5c which is mainly due to the close packing of UbcH5c-BRCA1 RING domain (as a result of covalent bonding). Since UbcH5c undergoes sliding motion on the NCP surface in addition to hinge motion, the energy of UbcH5c-BRCA1 has no obvious correlation with the UbcH5c(E2)-NCP distance change (correlation of the y axis with color scale shown in Figure S15a), thus showing no clear effect on UbcH5c upward motion. However, initially UbcH5c is oriented close to the NCP surface (E2-NCP distance is 12.83 Å), hence the repulsive forces corresponding to positively charged UbcH5c (net charge is +3.37 e) and H2A C-tail (net charge is +3.99 e) would result in a positive electrostatic energy between UbcH5c and H2A C-tail (the squared region in Figure S16a and the interaction energy during the initial 50 ns shown in Figures S15b-c). Consequently, this repulsion, together with some contribution from the BRCA1/BARD1 pull due to its tilt, creates an initial force on UbcH5c to pull it away from the surface.

To further explore how this repulsion caused by H2A C-terminal lysine residues promotes UbcH5c motility, BRCA1/BARD1-UbcH5c activity in the nucleosome when all five H2A C-terminal lysine residues are mutated to alanine (K486A, K487A, K493A, K495A and K497A) was tested. We performed 3 independent MD simulations of the mutated all-ALA systems, each for about 800 ns, where the only difference is the set of initial velocities. Interestingly, no uplift movement of UbcH5c was observed in any of the trajectories. The UbcH5c-NCP distance calculated and compared to the WT results in Figure 8a shows a pronounced peak around 23.5 Å for all-ALA mutation simulations indicating a steady distance between UbcH5c and NCP surface throughout the trajectories compared to the broad peak area of 2.1 - 67.8 Å obtained for WT representing the UbcH5c motions. For the WT trajectories, the two distinct broad peaks indicate the highest probability of finding UbcH5c around 17 Å and around 41 Å from the nucleosome (Figure 8a). Further, the large RMSF standard deviations of the H2A C-tail residues shown in Figures 8b and 8c suggest the structural mobility gained in all-ALA simulations in contrast to the WT.

**Figure 8:**
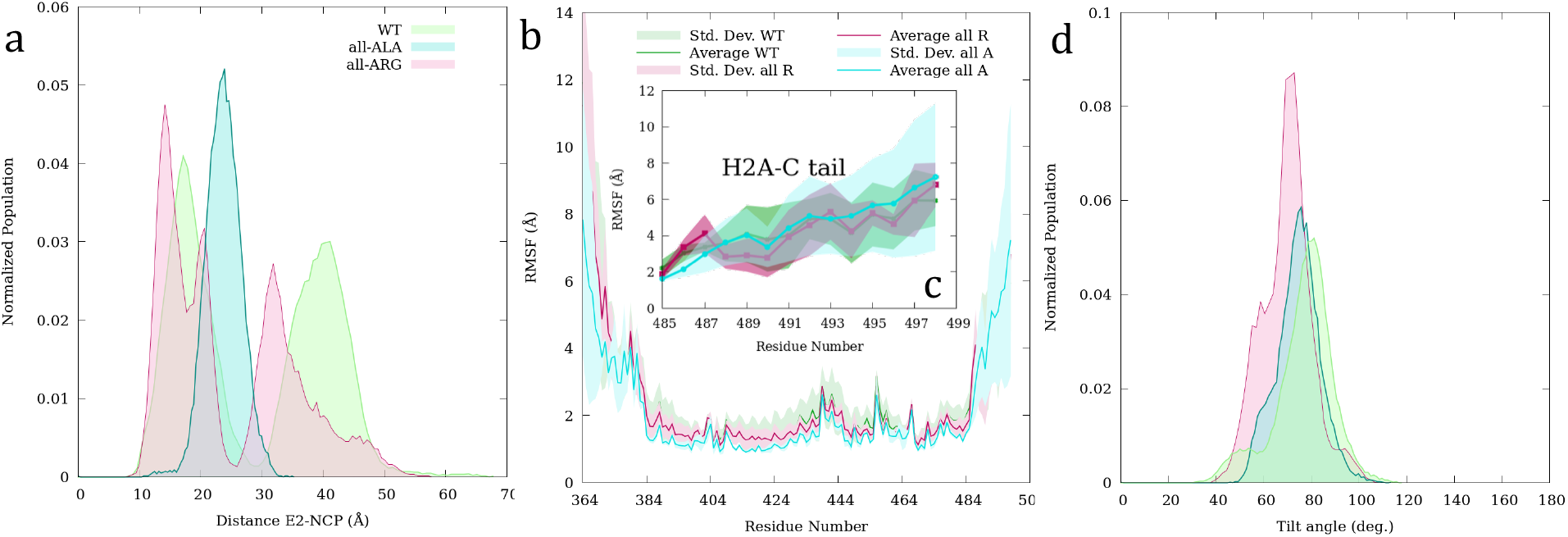
a. UbcH5c (E2)-NCP distance of WT, all alanine (all-ALA) and all arginine (all-ARG) systems. Distance is calculated between the residue D242 (C_*α*_) on UbcH5c and residue 1443 (P atom) on DNA. b. RMSF of the entire H2A chain. c. RMSF of the H2A C-tail of WT, all-ALA and all-ARG systems. d. BRCA1-H2B tilt angle of WT, all-ALA and all-ARG systems.

Moreover, the electrostatic interaction energies between UbcH5c and the H2A C-tail, DNA, BRCA1, H2B and BARD1 were compared with WT in Figure S16 (a-b). In all-ALA simulations, neither UbcH5c-H2A C-tail nor UbcH5c-DNA repulsive interactions were detected (squared regions in Figure S16b). Additionally, the DNA-UbcH5c attractive interactions are highly crowded around -142.2 kcal/mol in all-ALA system (in WT it is more crowded around -48.5 kcal/mol) due to the lack of electrostatic repulsion coming from the H2A C-tail on UbcH5c. Furthermore, the smaller BRCA1-UbcH5c interaction (highest population around -98.1 kcal/mol) compared to WT (highest population around -250 kcal/mol) and the distance between UbcH5c and NCP indicate the absence of UbcH5c hinge or sliding motion in all-ALA simulations. Taken together, the results explain how the lack of electrostatic repulsion in all-ALA mutated H2A C-tails affects the initial uplifting motion of UbcH5c in contrast to WT.

To further confirm the electrostatic effect stemming from the H2A C-tail to UbcH5c movements, three independent MD simulations of BRCA1/BARD1-UbcH5c with NCP were conducted mutating all five H2A C-tail lysine to arginine (all-ARG). As expected, UbcH5c undergoes a hinging motion perpendicular and a sliding motion parallel to the nucleosome surface in all-ARG simulations similar to WT. The UbcH5c-NCP distance in Figure 8a also shows two highly populated regions (shown in red) for all-ARG simulations confirming the uplifting behavior of UbcH5c similar to WT. This further establishes the importance of H2A C-tail electrostatics for the upward movement of UbcH5c from the surface.

Both arginine and lysine are positively charged basic amino acids with high aqueous pK_*a*_’s (∼10.5 for Lys and ∼12.5 for Arg) ^63^. The guanidine group of arginine allows interactions in three possible directions, which enables arginine to form more electrostatic interactions than lysine, and its higher pK_*a*_ can also generate more stable ionic interactions than lysine^63^. Thus, in all-ARG simulations, all H2A C-tail residues bind strongly to DNA (Figure S17), which diminishes their ability to reach the active site of UbcH5c, thereby inhibiting the potential for ubiquitination. As a result, experimental mutagenesis studies of H2A lysine to arginine show no/low ubiquitination efficiency compared to WT.^12,17,19^ Furthermore, BRCA1/BARD1 tilt angle results presented in Figure 8d and RMSF data shown in Figure S18 for WT, all-ALA and all-ARG simulations suggest that the H2A C-tail interaction has no direct effect on BRCA1/BARD1 motion on the NCP surface. Overall, our findings imply that UbcH5c is stiffer at the NCP surface in the absence of electrostatic repulsion from the H2A C-tail.

### Ubiquitination mechanism

The mechanism by which BRCA1/BARD1-UbcH5c ubiquitinates the lysine substrates of H2A C-tail has not been fully unraveled. In the BRCA1/BARD1-UbcH5c/NCP cryo-EM structure, the C_*α*_-C_*α*_ distance between the active site of UbcH5c and the K486 residue of the H2A ordered region is higher (∼ 19 Å) compared to that in Ring1b-bound UbcH5c structure (∼ 9 Å). Based on these distances, previous studies^12,19^ proposed that there is a seesaw-like motion for BRCA1/BARD1-UbcH5c that places the active site of UbcH5c away from the ordered C-tail region of H2A, and this position disfavors the ubiquitination of K486 and K487 in the H2A C-tail, but favors the ubiquitination of last three disordered C-tail lysine residues (K493, K495, and K497). In the current study, to provide an adequate molecular-level picture of the mechanism by which BRCA1/BARD1-UbcH5c ubiquitinates H2A C-tail lysine, MD-simulation trajectories were examined and the results as a function of time were taken into account. In WT MD run 1, which was simulated for 1050 ns, all major motions were observed for considerable time intervals. Therefore, to demonstrate the mechanism with time, the results of WT MD run 1 were accommodated.

In the previous sections, we identified the large amplitude upward/downward tilting movement of BRCA1/BARD1 and hinging movement of UbcH5c on the NCP surface. The comparison of BRCA1/BARD1 tilt angle results presented in Figures S11a and b with the changing C_*α*_-C_*α*_ distance between the active site residue K197 of UbcH5c and the H2A Ctail lysine residues presented in Figure S11c for WT MD run 1 as a function of time, clearly pronounces a relationship between these two movements. The BRCA1/BARD1-UbcH5c shows a see-saw dynamics on the NCP surface where BRCA1/BARD1 tilt angle decreases the UbcH5c distance increases and vice versa (Figure S11) confirming the proposed mechanism in the literature. PCA mode F and the MD trajectory in Video S2 further support these dynamics of BRCA1/BARD1-UbcH5c on the surface. Consistently, when all WT simulation data are considered, the UbcH5c-NCP distance and the BRCA1-H2B tilt angle in Figure S19 shows a negative correlation (when the distance decreases from 55 Å to 15 Å the angle increases from 30° to 94°) with some outliers due to other simultaneous movements (such as sliding, rotation, Tili-2).

Moreover, the trajectory snapshots shown in Figure 9 illustrate that when the hinging/sliding movements of UbcH5c distance the active site away from the surface of NCP (Figure 9b), the flexible disordered C-terminal tail of H2A also emanates from the surface to interact with UbcH5c and bring it back to the surface (Figure 9c). During this process, H2A C-tail lysine residues, such as K497 and K495, show the potential to approach the active site. The C_*α*_-C_*α*_ distance profiles for the UbcH5c active site (K197) and H2A C-tail lysine shown in Figures 6b (the closest distance traveled by residues K495 and K497 is approximately 12 Å) and S11c (for WT run 1) further confirm the likelihood of these residues reaching the active site. The interaction energies of the K495 and K497 residues of H2A with the K197 active site shown in Figure S11d exhibit strong repulsive interaction energies during 150–300 ns, supporting the mechanism. Additionally, the electrostatic interaction energy between K497 and the DNA plotted in Figure S11e further confirms the uplifting behavior of the disordered H2A C-tail towards UbcH5c. Initially, the K497 terminal lysine interacts strongly with DNA (energy is about -350 kcal/mol), where the K497 side chain is located within the DNA pocket. Once K497 escapes from the pocket, the interaction energy is reduced by 300 kcal/mol (Figure S11e). Over time (within 50-300 ns), the K497-DNA interaction increases (up to -325 kcal/mol), indicating that H2A tail returns to the NCP surface (Figure S11e). The MD simulations show that, during the above time period, sometimes K497 helps to bring UbcH5c to the surface by strongly interacting with oppositely charged residues in UbcH5c and/or approach to the active site of UbcH5c to potentially ubiquitinate (Figure 9).

**Figure 9:**
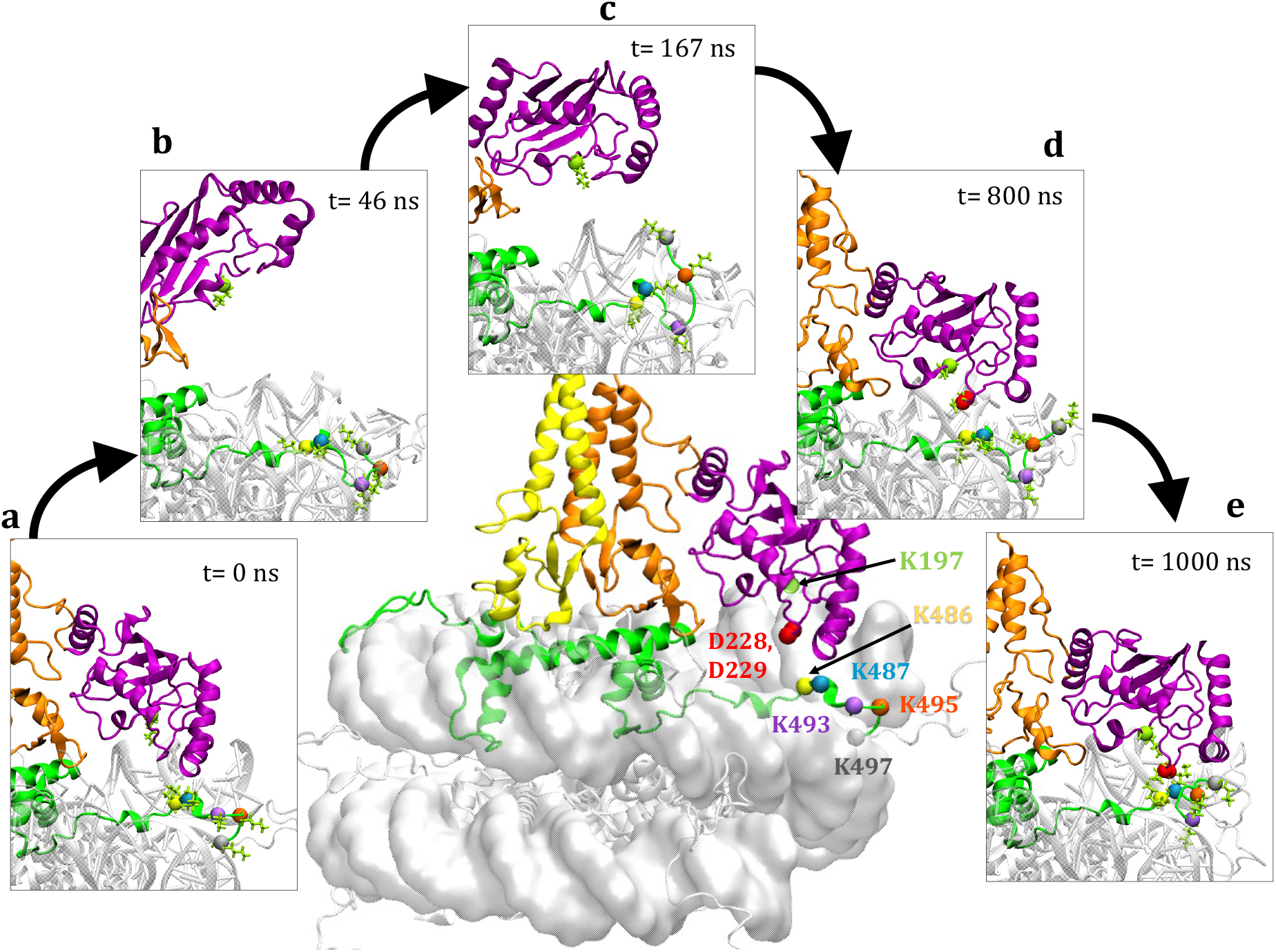
The mechanism showing the emanation of the C-terminal tail of H2A from the nucleosome surface to interact with UbcH5c active site (K197). Coloring as follows: DNA (gray), BARD1 (yellow), BRCA1 (orange), UbcH5c (purple), and H2A (green).

Although the disordered C-tail of H2A is free to leave the DNA site during this process, K493 in H2A disordered region remains trapped within a DNA-binding pocket (Figure 9). Figure S11c shows that the K197-K493 distance remains the highest compared to others, while in Figure S11e, the DNA-K493 energy remains around -130 kcal/mol throughout the simulation. Figure 7 also shows a comparatively low contact frequency of K493 with UbcH5c, and zero/none interaction energy between K493-K197 is obtained in Figure S11d. These results suggest that residue K493 acts as an anchor, thus its arrangement results in limited flexibility in the H2A C-tail. This K493 anchoring behavior was confirmed in all WT trajectory results. The change in distance (Δd) between the five lysine residues in the H2A C-tail relative to one of the selected DNA locations (residue 1443) depicted in Figure S20a shows the highest Δd distribution around 5 Å for K493, while K495 and K497 show the highest population around 14 Å and 19 Å, respectively. The normalized electrostatic interaction energy distributions of the five lysine residues in the H2A C-tail with the K197 active site of UbcH5c depicted in Figure S20b for all WT trajectories also show the weakest interactions between K493 and K197. Supporting our results, Kalb *et al*.^17^ reported the ubiquitination of only disordered H2A C-tail lysine 495 and 497, but not lysine 493.

In addition, our results show that when UbcH5c returns to the nucleosome surface, K497 moves away from the DNA end (Figures 9d and S11c) to position the UbcH5c active site closer to other lysine residues to transiently transfer Ub to those Ub acceptor lysines on the H2A C-tail. The loop region of UbcH5c containing residues D228 and D229 shows a significant contribution to bring K197 closer to the lysine residues on the H2A C-tail, once UbcH5c returns to the NCP surface (Figure 9e). The electrostatic interaction energies shown in Figures S21a and S21b confirm that residues D229 and D228 strongly interact with K197 and H2A C-terminal lysine residues and act as intermediaries to bring the residues closer together. Figures S21c and S21d show the individual electrostatic interaction energies between D228/D229 and H2A C-tail lysine residues indicating strong K486 attractions with D228/D229. Meanwhile, Figures S20b and 9e state that K487 gets closer to the UbcH5c active site compared to others. This implies that D228/D229 indirectly helps to bring the active site closer to the H2A lysines. Consequently, all 486-497 lysine residues in the H2A C-tail show the ability to be ubiquitinated by BRCA1/BARD1 in the presence of UbcH5c, although K487 shows the highest probability of being ubiquitinated once UbcH5c is positioned closer to the nucleosome surface (Figure 9). Contact frequency results from WT simulations also show the highest probability for K487 and the lowest probability for K493 (Figure 7).

### K487 on the H2A C-tail shows the highest probability of ubiquitination

To attain a quantitative picture of how the location of lysines in the H2A C-tail determines the efficiency of ubiquitination by BRCA1/BARD1-UbcH5c, each lysine in the H2A tail was mutated to an alanine. Five different MD simulations of K486A, K487A, K493A, K495A and K497A were performed and the probability of ubiquitination of each lysine was further investigated. Each simulation was conducted for 500 ns and subjected to various analyses, including RMSF, interaction energy, and distance calculation to investigate the effect of mutation on the structure and dynamics of H2A and UbcH5c.

Comparing the RMSF results in Figures 8b and 10a, it was found that a single lysine mutation does not confer more flexibility and mobility to the H2A C-tail in contrast to all-ALA mutations. However, in the K497A mutant simulation, the end of the H2A disordered C-tails gained more mobility compared to WT due to the lack of electrostatic interactions with DNA (Fig. 10b). Interestingly, the K487A mutated system reports the highest RMSF of the H2A C-tail compared to others (Figure 10b). This indicates that the loss of electrostatic force exerted by K487 results in greater flexibility for the rest of the C-tail. It also suggests an important contribution of H2A K487 to the dynamics of the H2A C-tail. Comparison of time-averaged total non-bonded interaction energies between the active site UbcH5c (K197) and H2A C-tail residues (486, 487, 493, 495, and 497) in all single mutant and WT simulations in Figure 10c indicate that K487 shows the highest probability of reaching the active site (the more repulsive the energy, the closer it is). When K487 is mutated, K486 and K493 get closer to the active site (Figure 10c). Taken together, the results uncover that K487 in the H2A C-tail reaches the closet to the UbcH5c active site; implying the highest probability of ubiquitination by BRCA1/BARD1-UbcH5c.

**Figure 10:**
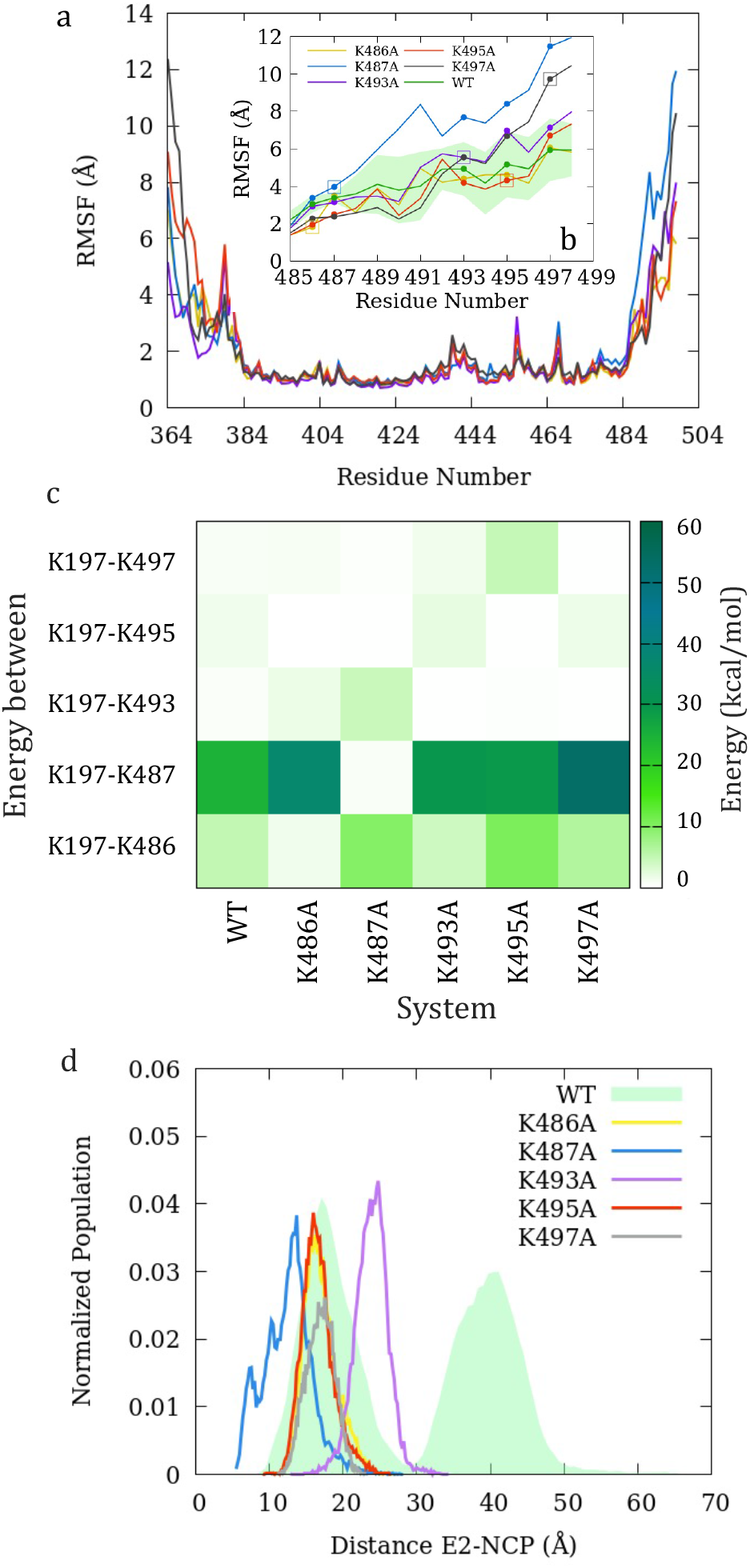
a. RMSF of H2A in all single mutants and WT simulations. b. Inset shows the RMSF of H2A C-tail. WT results are shown in green with standard deviation. All the mutated residues are squared. c. Total non-bonded interaction energy between UbcH5c active site (K197) and five lysine residues in H2A C-tail. d. Normalized distribution of UbcH5c (E2)-NCP distance (distance between the residue D242 (C_*α*_) on UbcH5c and 1443 (P atom) on DNA) for all WT, and single mutant trajectories.

Meanwhile, the highest interaction of K497 (last lysine residue in the disordered H2A tail) with the active site of UbcH5c was obtained in the K493A and K495A simulations (Figure 10c) because in the absence of the electrostatic effects of K493 and K495, the disordered H2A C-tail gains more mobility which increases the ability of K497 to reach the active site. On average, across all simulations, K493 shows the weakest interactions with the UbcH5c active site (Figure 10c), confirming its anchoring behavior. Further, when K493 mutates to alanine, the interactions between the active site of UbcH5c with K497 and K495 increase due to the lack of the anchoring effect of K493 (Figure 10c). Altogether, our MD simulations predict that K487 is most likely to reach the active site of UbcH5c and thus shows the highest probability of being ubiquitinated by BRCA1/BARD1-UbcH5c, while other H2A C-tail lysine residues also show some potential for ubiquitination. According to Kalb *et al*.^17^ K495-K497 and the more common K487 ubiquitination perform very similar functions in chromatin. Interestingly, in contrast to WT simulations, little or no UbcH5c hinging/sliding motions were observed (Figure 10d) in all single mutant simulations, further confirming the importance of electrostatic effects stemming from the 5 lysine residues for the dynamics of UbcH5c on the NCP surface.

## CONCLUSIONS

BRCA1/BARD1-dependent ubiquitination of the nucleosomal H2A C-terminal tail is critical for its function as a tumor suppressor and has attracted considerable attention in the recent literature. The current study provides insight into the mechanism by which BRCA1/BARD1UbcH5c ubiquitinates H2A C-tail lysine residues using MD simulations. Simulations results show that the BRCA1 (R73) and BARD1 (W329) nucleosome-binding factors confine them to the NCP surface and form a unique pivot-like architecture that allows characteristic movements of the BRCA1/BARD1-UbcH5c complex on the surface. PCA reveals the flexible states accessible to BRCA1/BARD1 and UbcH5c and shows that BARD1 and BRCA1 can undergo two tilting movements (Tilt-1 and Til-2), N-terminal movement (N-tail motion) and RING translation motion while UbcH5c can undergo two large-scale hinging (perpendicular to the nucleosome surface) and sliding (parallel to the nucleosome surface) movements and a small-amplitude rotational motion along the principal axis, passes through the protein. The electrostatic interactions between BRCA1/BARD1 with H2B/H2A and UbcH5c play an important role in determining the movement of BRCA1/BARD1, while the electrostatic interactions between UbcH5c with H2A C-tail and DNA primarily affect the structural dynamics of UbcH5c on the surface.

Comparison of WT with all-ALA and all-ARG mutants suggests that the electrostatic repulsion of UbcH5c by the H2A C-tail lysine drives the initial hinging/sliding motion of UbcH5c. As UbcH5c moves away from the surface, its rotational motion causes some charged residues to face the NCP surface and strongly interact with oppositely charged residues on the H2A C-tail and DNA. This arrangement allows the flexible H2A disordered C-tail to leave the NCP surface and interact with UbcH5c, bringing it back to the surface with the help of the K493 anchoring effect. The results predict different scenarios in which residues K495 and K497 in the disordered H2A C-tail show the possibility of being ubiquitinated when emanating away from the NCP surface, whereas all the lysine residues of the H2A Ctail show the possibility of ubiquitination when UbcH5c returns to the nucleosome surface. The loop region of UbcH5c containing residues D228 and D229 was shown to significantly contribute to bring the active site closer to the lysine residues in the H2A C-tail once UbcH5c returned to the NCP surface.

The results obtained from five single mutant simulations (K486A, K487A, K493A, K495A, and K497A) predict the probability of the H2A C-terminal lysine reaching the active site, thus, the effect of lysine location on ubiquitination efficiency as the order of K487 > K486 > K497 > K495 >> K493. Altogether, our findings suggest that the dynamics of the H2A Cterminal tail, as well as the highly mobile UbcH5c, define ubiquitination capacity. Therefore, within a ubiquitination-capable framework, the UbcH5c can transiently transfer Ub to lysine targets on the H2A C-tail. In the next step, we will investigate the role of Ub in the ubiquitination of H2A, which we expect to facilitate tilting/hinging/sliding motions due to steric restrictions. Finally, our results reveal the importance of histone-derived electrostatic effects for BRCA1/BARD1-UbcH5c dynamics on the surface of NCPs that regulate the ubiquitination machinery. The findings of this study have provided unrevealed insights into the mechanism of H2A C-tail ubiquitination and the electrostatic interactions between PTM elements and histone tails, which will aid in controlling the communication between these elements for the design of potential therapeutics in the future.

## Author Contributions

H.T. designed and supervised the project; D.T.S.R. performed the computational studies and analyses; H.T. and D.T.S.R. analyzed the data and wrote the manuscript together.

## Conflicts of interest

There are no conflicts to declare.

## Acknowledgements

Computational resources for this research were provided by High Performance Computing center at The University of Texas at Dallas and Texas Advanced Computing Center (TACC).

## Notes

### Competing Interest Statement

The authors have declared no competing interest.

