## Supplementary material for "Histone Tail Electrostatics Modulate E2-E3 Enzyme Dynamics: A Gateway to Regulate Ubiquitination Machinery": Table S1

Table S1: The sequence identity scores obtained from homology modeling [a]

| Chain | # RMSD (Å) | Template modeling score | % sequence identity | % sequence similarity | Alignment coverage (%) |
|---|---|---|---|---|---|
| N (H2A) | 0.08 | 0.98 | 92.8 | 96.8 | 100.0 |
| n (H2A) | 0.07 | 0.98 | 92.8 | 96.8 | 100.0 |
| O (H2B) | 0.10 | 0.99 | 93.4 | 98.4 | 100.0 |
| o (H2B) | 0.09 | 0.99 | 93.4 | 98.4 | 100.0 |
| P (H3.2) | 0.13 | 1.00 | 98.5 | 98.5 | 100.0 |
| p (H3.2) | 0.08 | 1.00 | 98.5 | 98.5 | 100.0 |
| Q (H4) | 0.11 | 1.00 | 100.0 | 100.0 | 100.0 |
| q (H4) | 0.08 | 1.00 | 100.0 | 100.0 | 100.0 |
| DNA | 1.39 | | | | |

[a] Template modeling score = 1 describes a perfect match.

Table S2: Total simulation time

| System | Trial No | Production time (ns) |
|---|---|---|
| WT | 1 | 1050 |
| WT | 2 | 500 |
| WT | 3 | 1100 |
| WT | 4 | 500 |
| WT | 5 | 500 |
| K all R | 1 | 1000 |
| K all R | 2 | 500 |
| K all R | 3 | 500 |
| K all A | 1 | 800 |
| K all A | 2 | 800 |
| K all A | 3 | 800 |
| K486A | 1 | 500 |
| K487A | 1 | 500 |
| K493A | 1 | 500 |
| K495A | 1 | 500 |
| K497A | 1 | 500 |
| Total time (ns) | | 10550 |

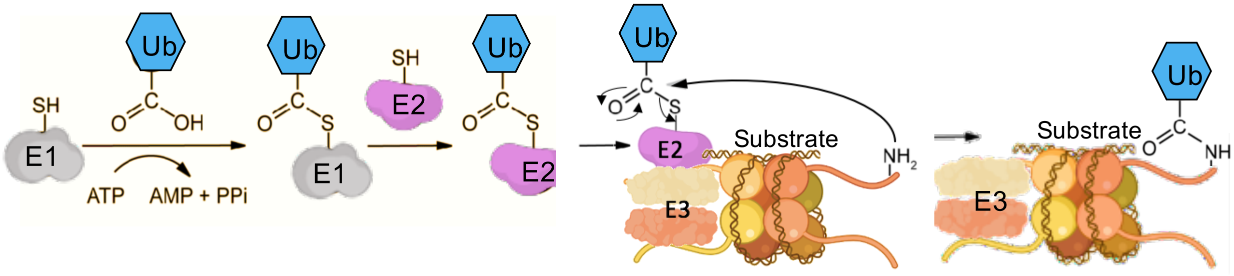


Figure S1: Ubiqutination mechanism.

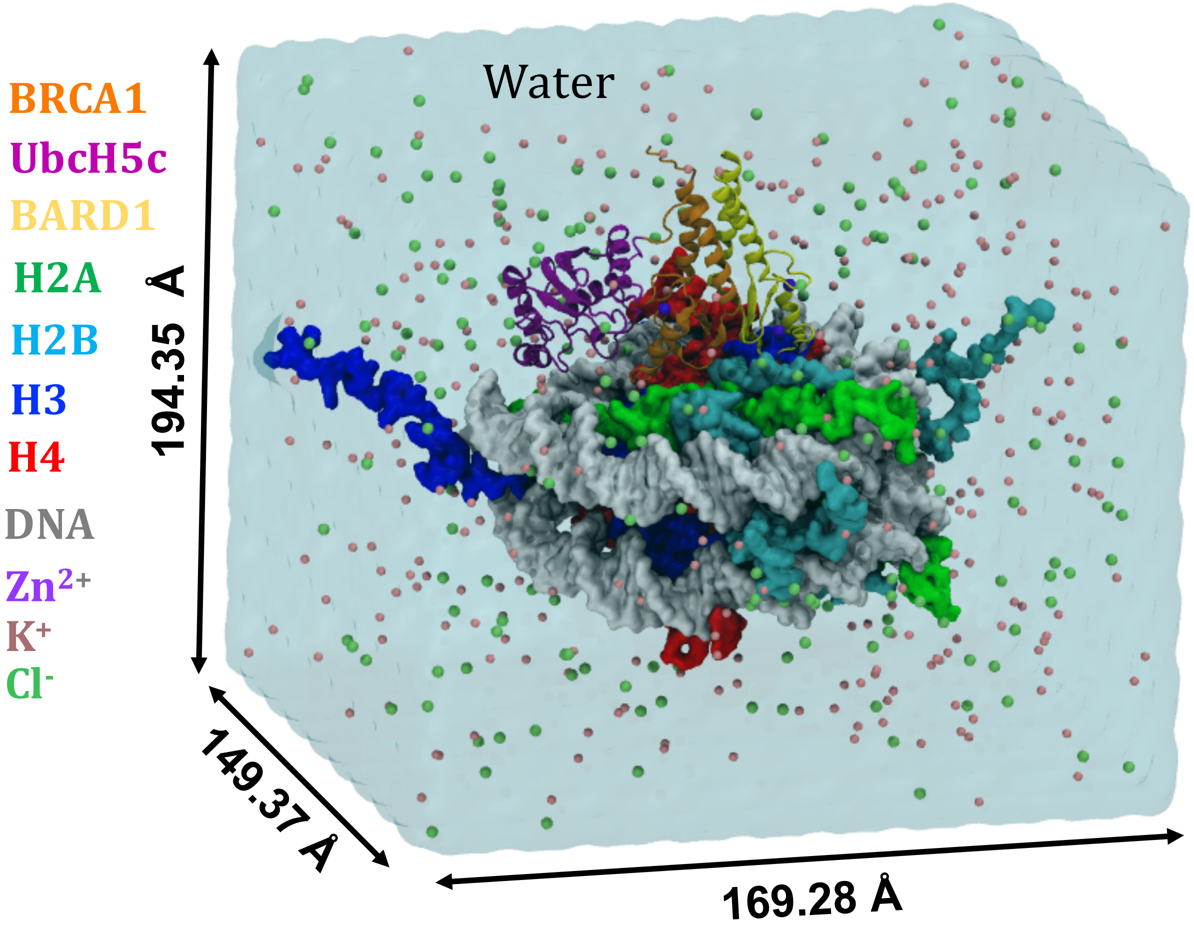


Figure S2: The MD simulation setup.

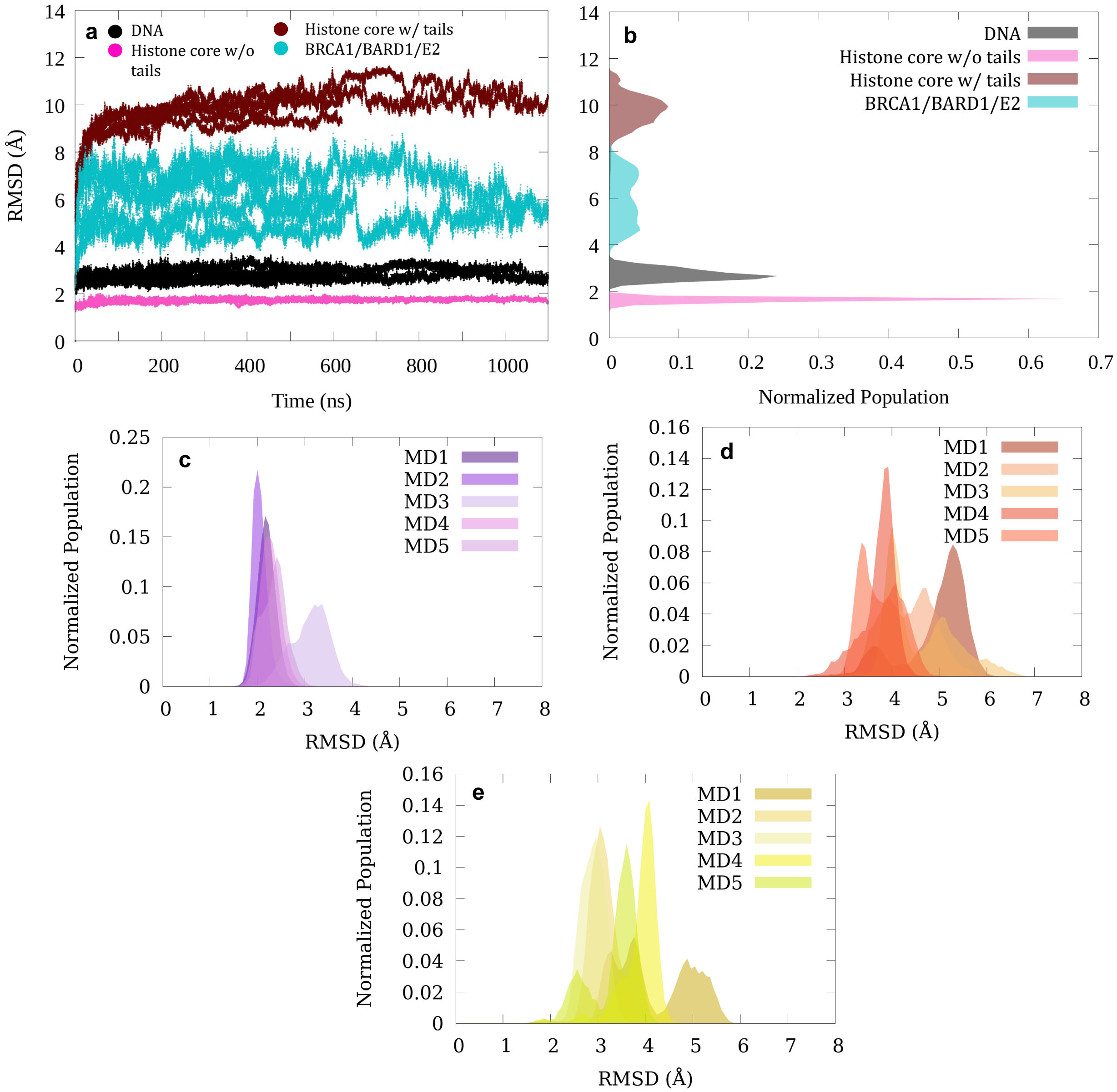


Figure S3: a. Backbone root mean square deviations (RMSDs) of DNA, BRCA1/BARD1-UbcH5c and nucleosome core which contains only 8 histone proteins with (Histone core w/ tails) and without (Histone core w/o tails) histone tails of all WT simulations with respect to the initial conformation. b. Normalized RMSD populations of DNA, histone core without tails, histone core with tails, and BRCA1/BARD1-UbcH5c domain in all WT simulations. Normalized RMSD populations of c. UbcH5c, d. BRCA1, and e. BARD1 domains in 5 different WT MD runs with respect to the initial configuration.

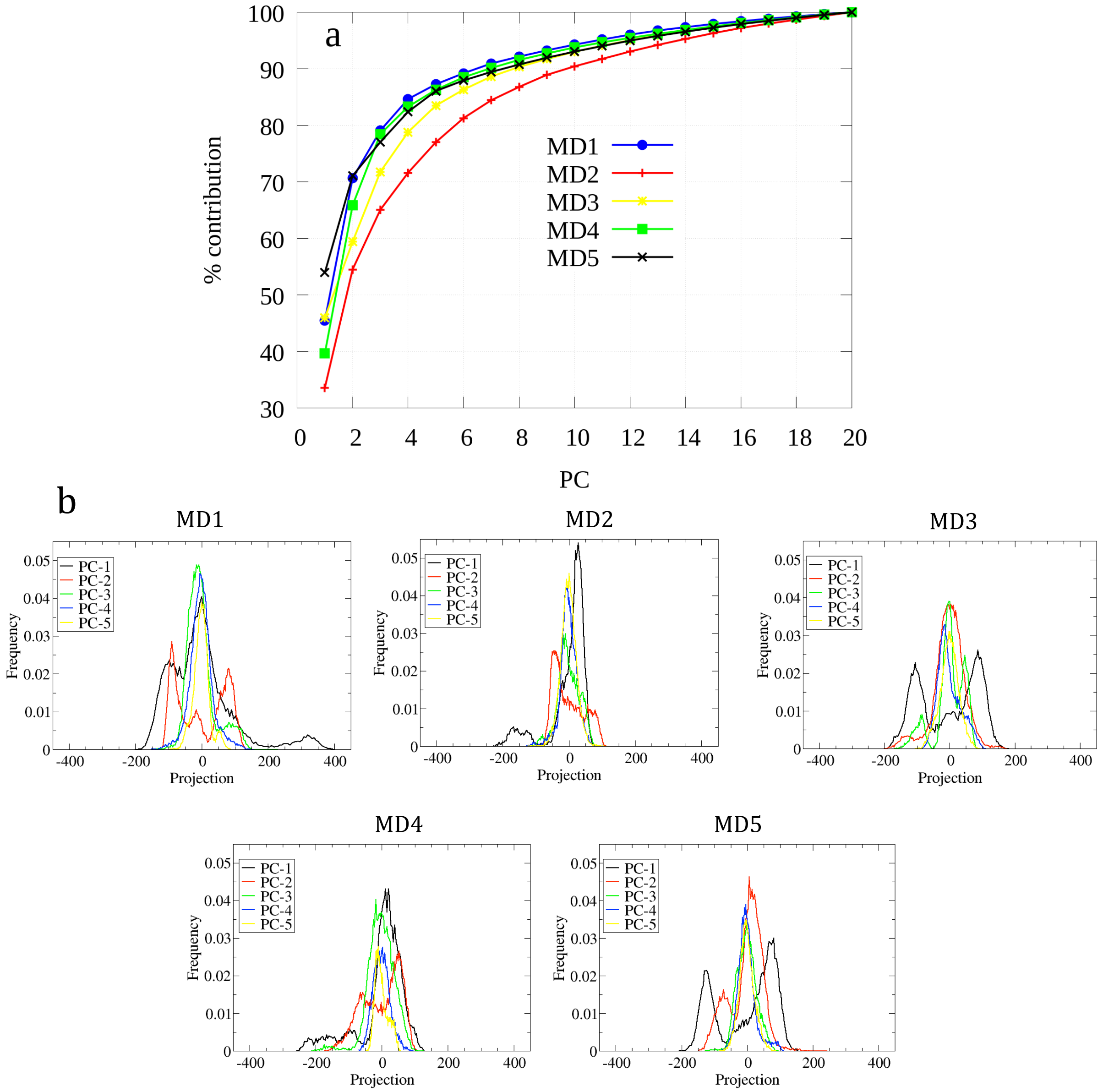


Figure S4: a. The cumulative contribution to variance in the BRCA1/BARD1-UbcH5c principal components obtained for all WT simulations. For the eigenvectors obtained from principal component analysis of five independent WT MD simulations, individual contributions to conformational changes are given. b. Normalized histograms of the calculated projections of first 5 PCs of different WT MD runs.

| Mode | | | MD1 | MD2 | MD3 | MD4 | MD5 | Total occurrence |
|---|---|---|---|---|---|---|---|---|
| A | 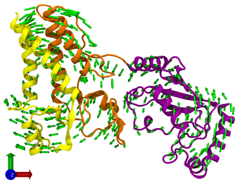 | 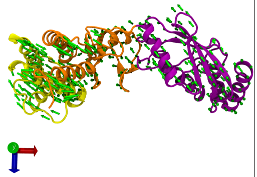 | PC1 45% | PC4 6% | PC1 34% | PC3 12% | PC1 40% | 137% |
| B | 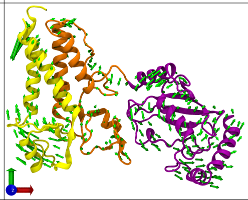 | 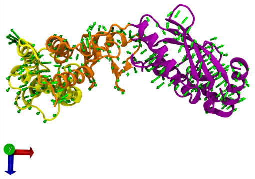 | PC2 25% | PC1 33% | PC2 19% | PC1 44% | PC3 8% | 128% |
| C | 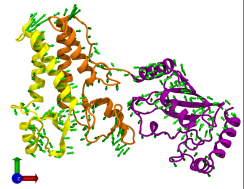 | 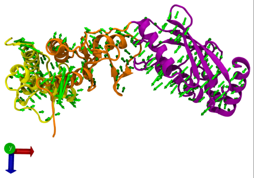 | PC3 8% | PC2 20% | PC6 5% | PC4 7% | PC4 4% | 44% |
| D | 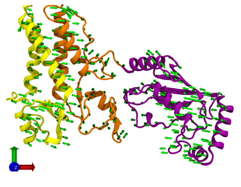 | 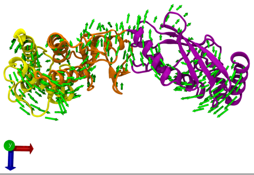 | PC4 6% | PC5 5% | PC4 6% | PC2 13% | PC2 26% | 56% |
| E | 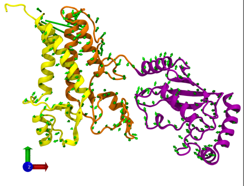 | 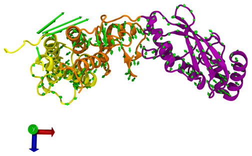 | PC5 3% | PC3 10% | PC3 10% | PC5 5% | PC6 4% | 32% |
| F | 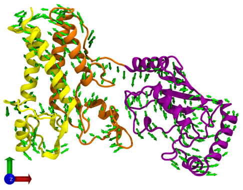 | 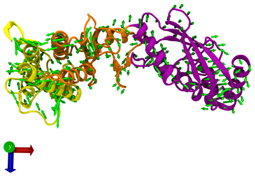 | PC6 3% | PC9 2% | PC7 4% | PC7 4% | PC5 4% | 17% |

Figure S5: Vector field representation of the 6 principal component modes (A-F) obtained from 5 different MD simulations for the entire BRCA1/BARD1-UbcH5c domain. For ease of visualization, only BRCA1/BARD1-UbcH5c are shown here. The NCP is positioned parallel to the XZ plane on the trajectories. The percent contribution of each mode to the variance in different MD runs is marked in blue (values were obtained from Figure S4a).

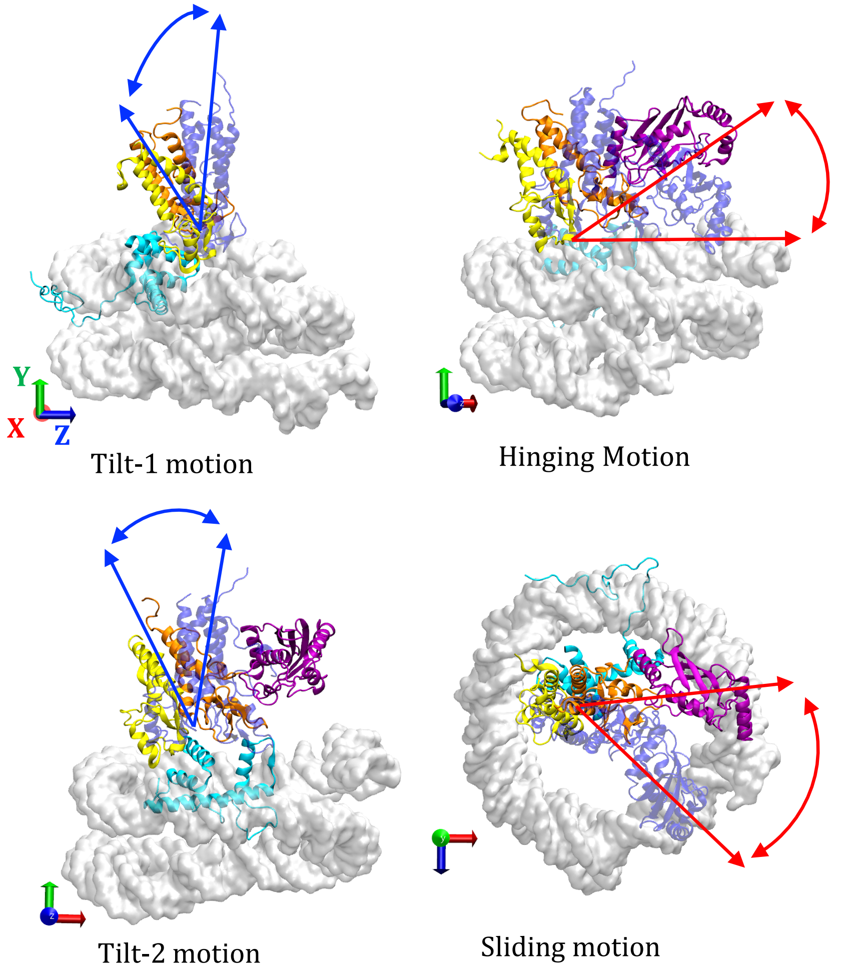


Figure S6: BRCA1/BARD1 and UbcH5c motions observed on the NCP surface compared to their initial location. For ease of visualization, only DNA, H2B and BRCA1/BARD1-UbcH5c were shown. Initial configurations are colored in transparent navy blue. In Tilt-1 motion BRCA1/BARD1 tilt is directed towards the H2B $\alpha$C helix (labeled in Figure S8a) and in Tilt-2 motion BRCA1/BARD1 tilts towards the YZ plane parallel to XY plane. In "hinging motion", UbcH5c displays a hinge around its interface with BRCA1. In "sliding motion", BRCA1-connected UbcH5c slides on the nucleosome surface parallel to the ZX plane. Coloring as follows: DNA (gray), BARD1 (yellow), BRCA1 (orange), UbcH5c (purple), and H2B (cyan).

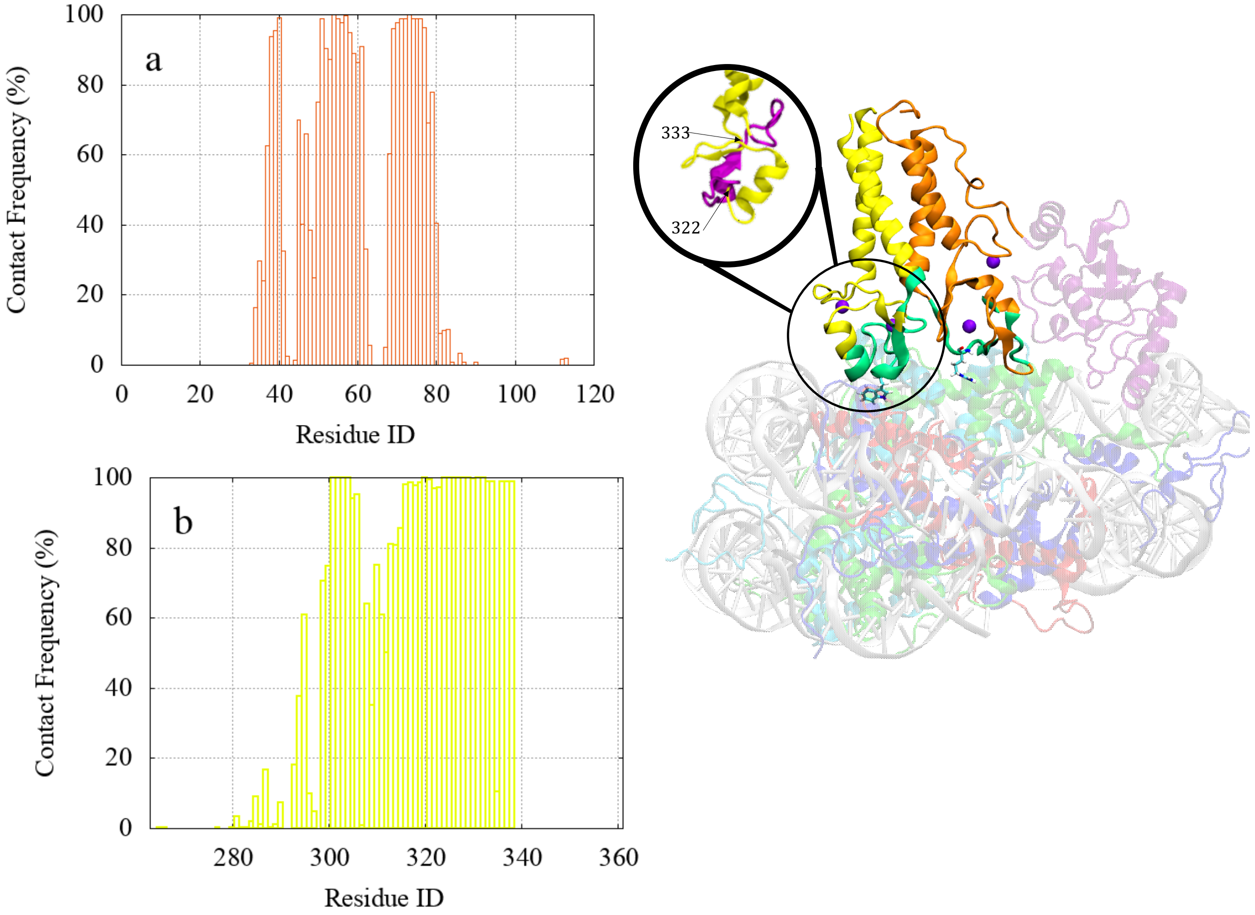


Figure S7: a. BRCA1 (orange) and b. BARD1 (yellow) contact frequency analysis. Regions with > 95% frequency are highlighted in the right panel (in mint green). Coloring as follows: DNA (gray), BARD1 (yellow), BRCA1 (orange), UbcH5c (purple), and H2B (cyan), H2A (green), H3 (blue), H4 (red). The enlarged panel of the BARD1 RING domain shows residues in the 322-333 region colored in magenta.

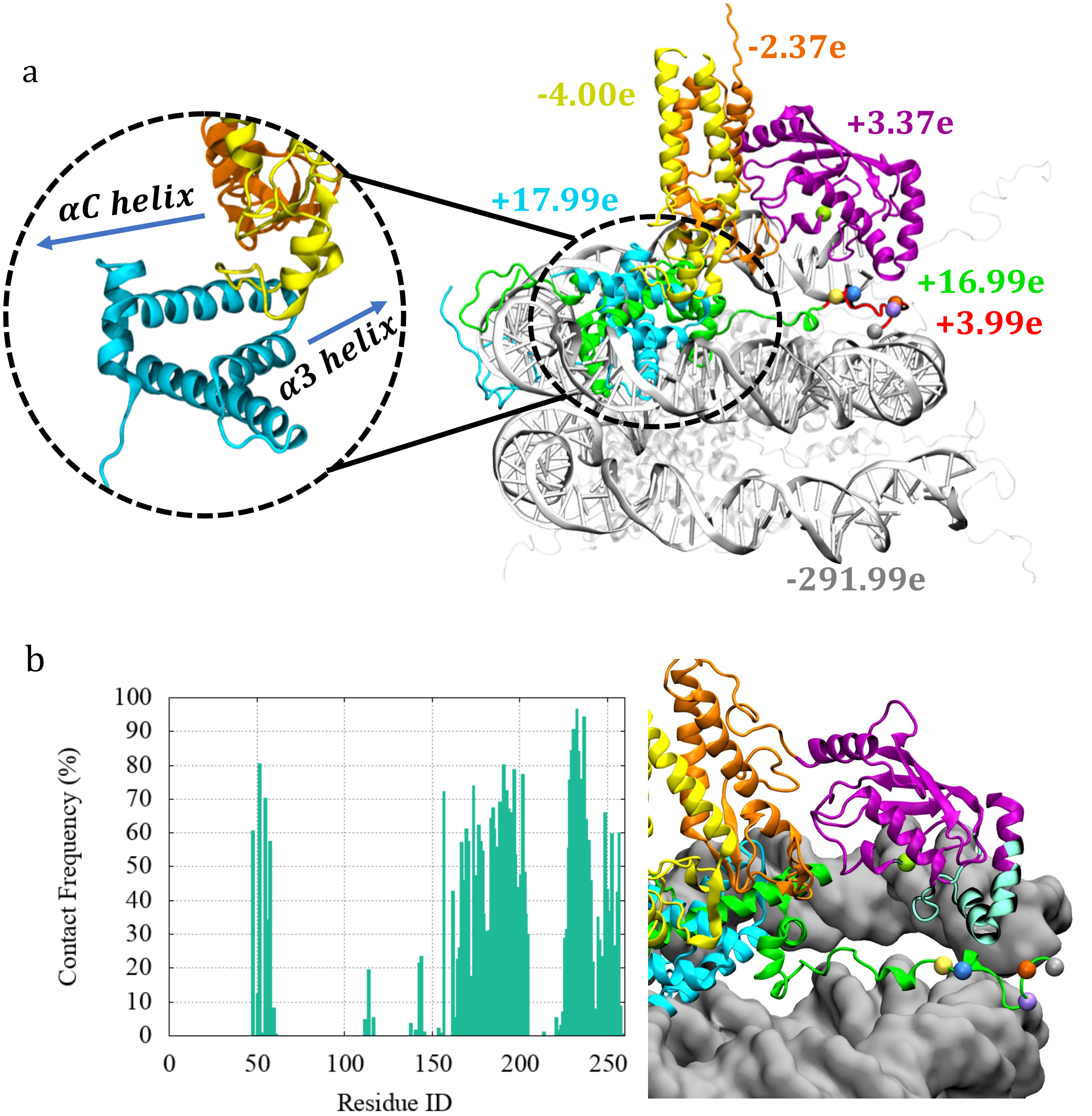


Figure S8: a. The right panel shows the net charges of different domains. $C_\alpha$ atoms of H2A C-tail lysines and UbcH5c active site are colored as follows: K197 (yellowish-green), K497 (gray), K495 (orange), K493 (purple), K487 (blue) and K486 (yellow). The left panel shows a zoomed-in view of a part of the histone H2B protein with the $\alpha$ helices labeled. b. Contact frequency results of BRCA1/UbcH5c - DNA for all WT simulations. Percentage time that residues in BRCA1/UbcH5c can be found close (<7 Å) to DNA. $C_\alpha$ of H2A C-tail lysines are shown in spheres. Residues K52, P230, L231, V232, P233 and R237 were found >80% of the time close to the DNA. Coloring as follows: DNA (gray), BARD1 (yellow), BRCA1 (orange), UbcH5c (purple), and H2B (cyan), H2A (green), H2A C-tail (red) and UbcH5c region that contacts H2A C-tail/DNA is colored in mint green.

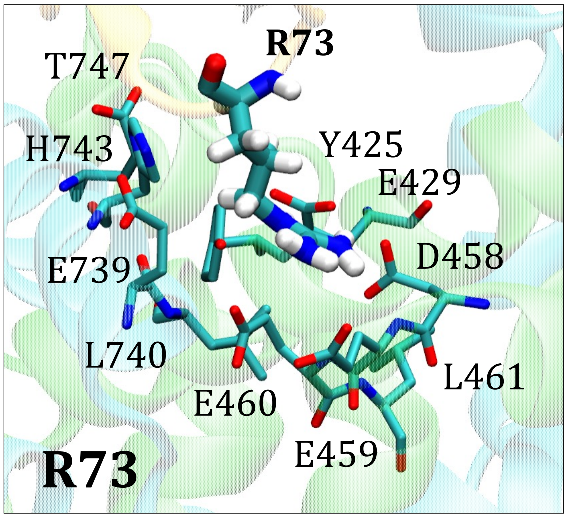


| Residues around R73 | H bond formation frequency (%) |
|---|---|
| T747 (H2B) | 5.000 |
| E739 (H2B) | 23.126 |
| Y425 (H2A) | 33.248 |
| E429 (H2A) | 99.103 |
| E460 (H2A) | 56.122 |
| E459 (H2A) | 38.820 |
| D458 (H2A) | 99.290 |
| L740 (H2B) | 8.118 |
| H743 (H2B) | 10.286 |
| L461 (H2A) | 23.632 |

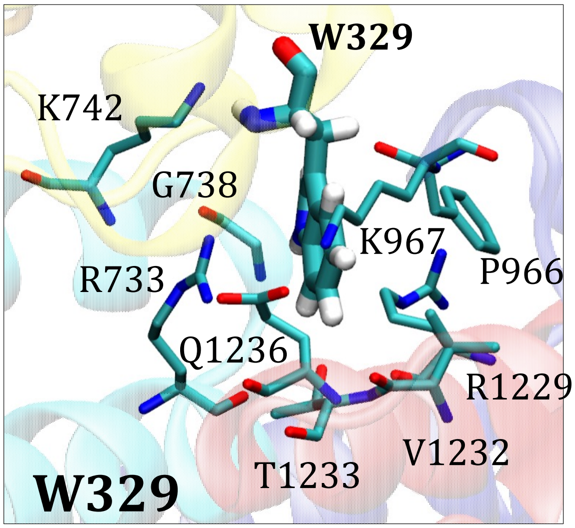


| Residues around W329 | H bond formation frequency (%) |
|---|---|
| K967 (H3) | 5.106 |
| R733 (H2B) | 40.118 |
| V1232 (H4) | 5.010 |
| Q1236 (H4) | 6.020 |
| P966 (H3) | 5.010 |
| G738 (H2B) | 6.016 |
| T1233 (H4) | 8.076 |
| K742 (H2B) | 15.612 |
| R1229 (H4) | 19.224 |

Figure S9: BRCA1 R73 and BARD1 W329 anchoring motifs and their surrounding residues (left). Hydrogen bond frequencies between the anchor residues and their surrounding residues (right). Coloring as follows: BARD1 (yellow), BRCA1 (orange), UbcH5c (purple), and H2B (cyan), H2A (green), H3 (blue), H4 (red), carbon (dark-cyan), oxygen (red), and nitrogen (blue).

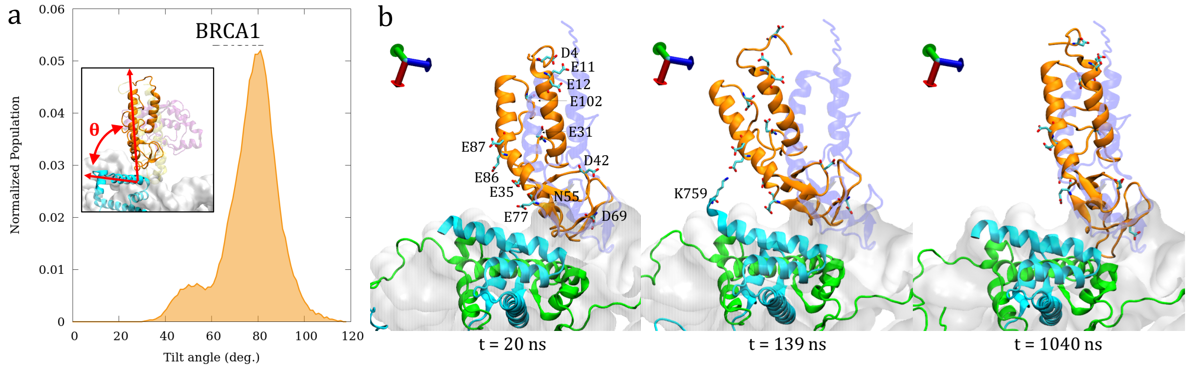


Figure S10: a. BRCA1 tilt angle distribution for WT trajectories. b. The directional Tilt-1 movement of BRCA1 relative to its initial configuration. Residues in BRCA1 contribute to the tilting motion (that show electrostatic energy higher than -100 kcal/mol with H2B and H2A in Figures S12b and S12d) are labeled. Coloring as follows: DNA (gray), BRCA1 (orange), H2B (cyan), H2A (green), initial structure (transparent blue), carbon atoms (dark cyan), oxygen (red), and nitrogen (dark blue).

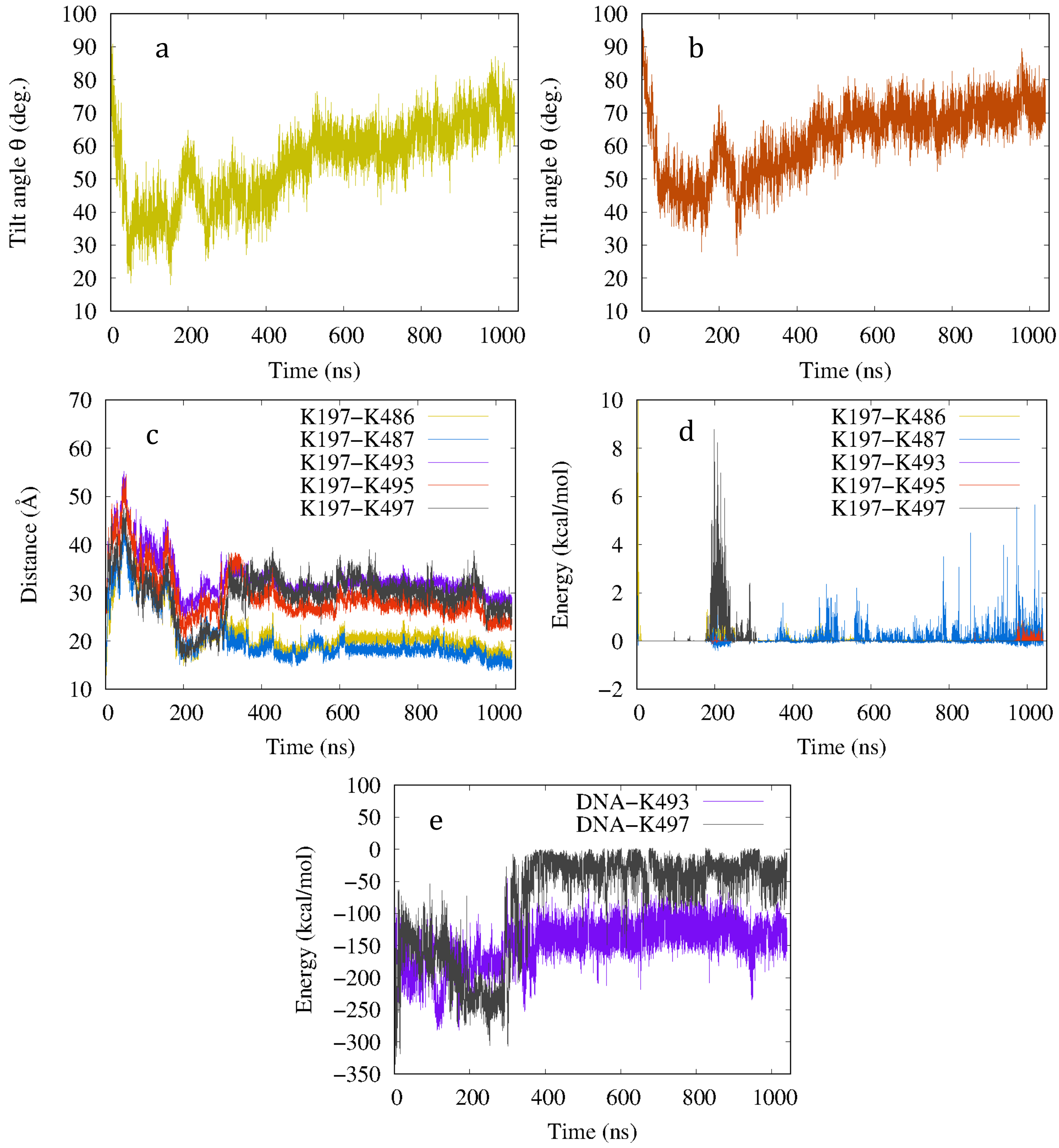


Figure S11: WT MD run 1 results. a. BARD1 and b. BRCA1 tilt angle change with time. c. Changing $C_\alpha$-$C_\alpha$ distance between K197 active site and H2A lysines with time. d. Interaction energy between 5 lysine residues in the H2A C-tail and active site K197 as a function of time. e. H2A C-terminal K493 and K497 residues interaction energy with DNA.

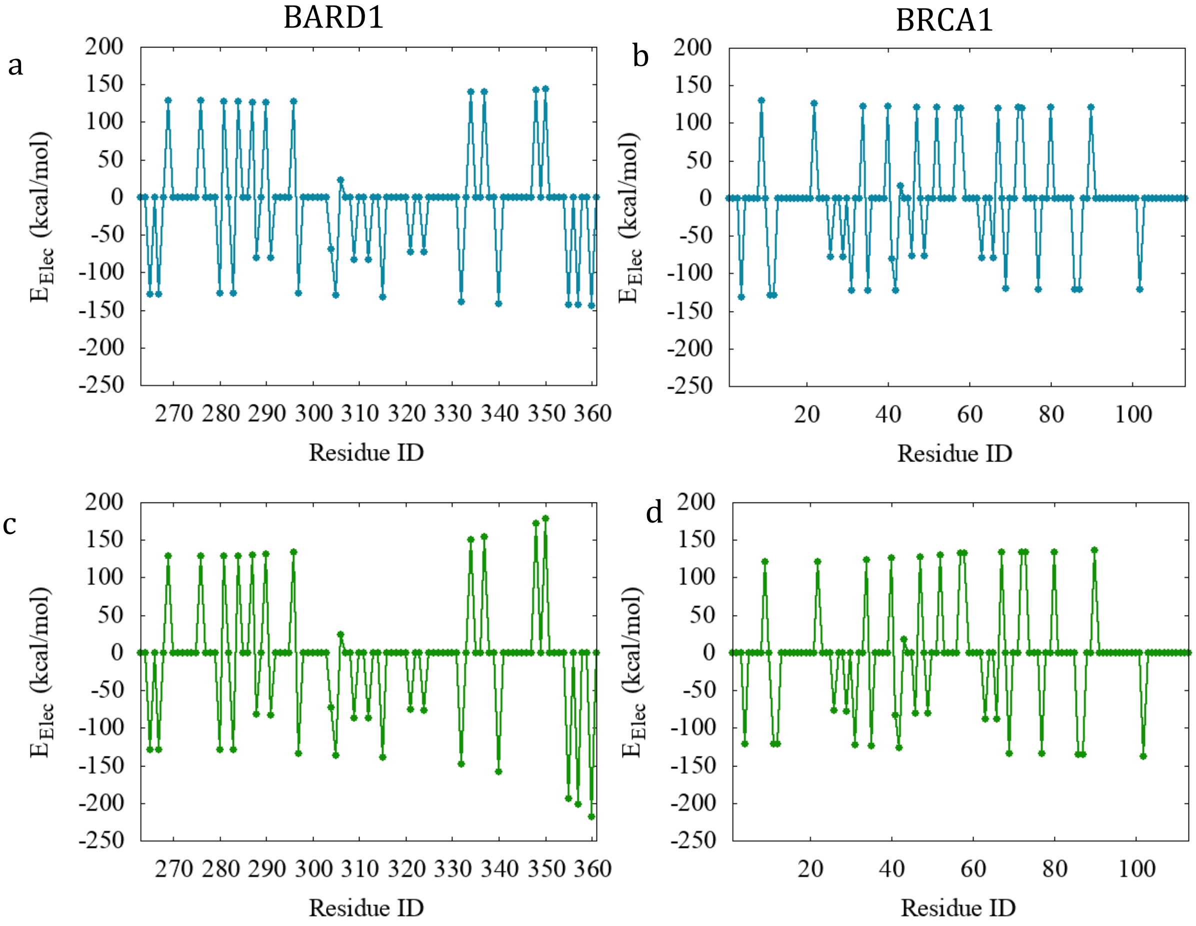


Figure S12: a. Average electrostatic interaction energies of residues on BARD1 with H2B for all WT simulations. b. Average electrostatic interaction energies of residues on BRCA1 with H2B for all WT simulations. c. Average electrostatic interaction energies of residues on BARD1 with H2A for all WT simulations. d. Average electrostatic interaction energies of residues on BRCA1 with H2A for all WT simulations. Results obtained from EDA.

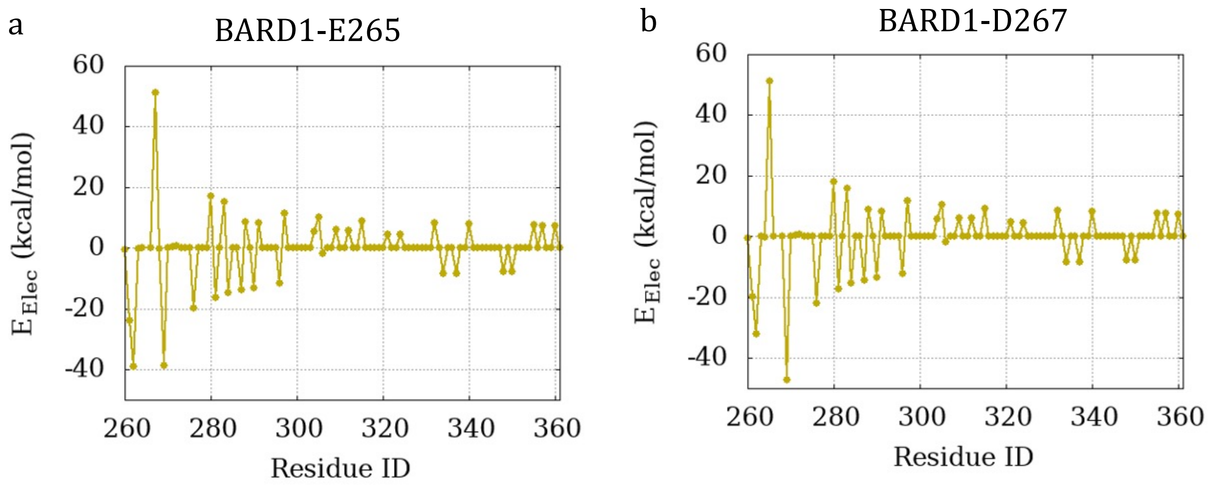


Figure S13: Average electrostatic interaction energy results for N-terminal E265 and D267 residues on BARD1 with the other residues on BARD1. Results obtained from EDA.

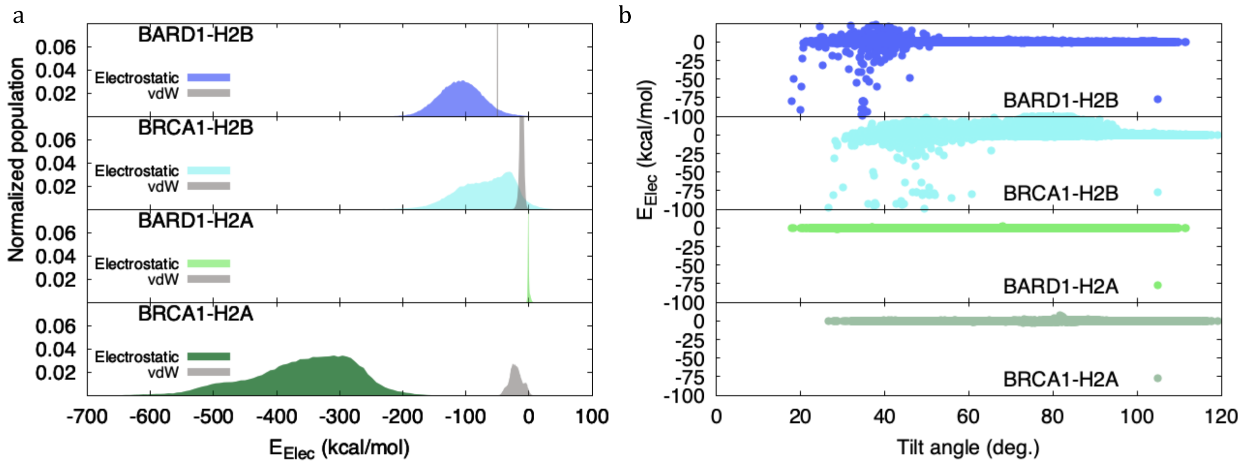


Figure S14: a. Electrostatic and vdW interaction energy between entire BRCA1 and BARD1 with H2B and H2A for WT. The BRCA1/BARD1 interaction energy with DNA is significantly smaller compared to BRCA1/BARD energy with H2B or H2A. b. Electrostatic energy between BRCA1 and BARD1 helical region with H2B and H2A compared to BRCA1 and BARD1 tilt angle.

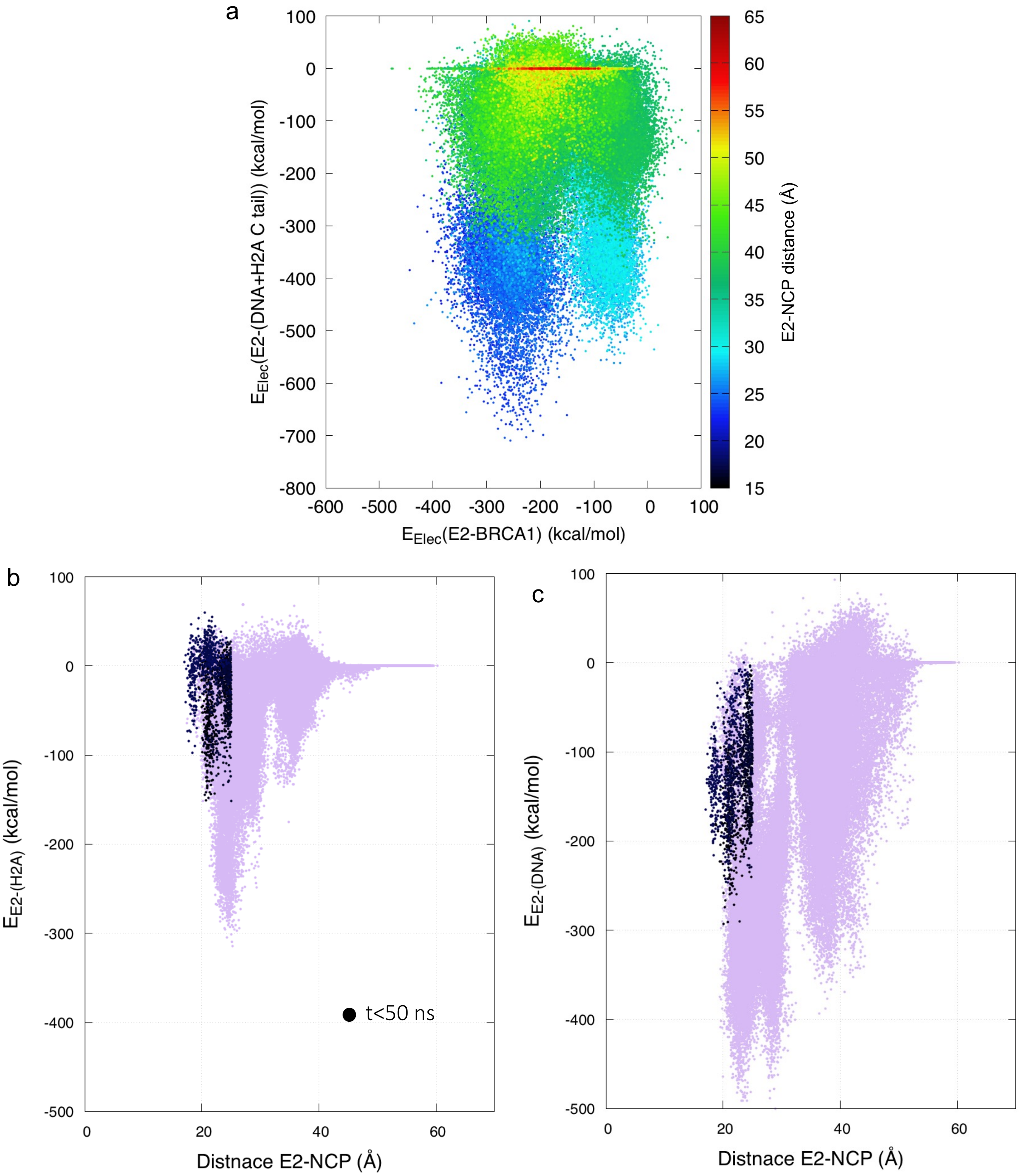


Figure S15: a. The electrostatic interaction energy between UbcH5c (E2) and (DNA+H2A C-tail) versus electrostatic interaction energy between UbcH5c and BRCA1. Colored as a function of UbcH5c-NCP distance. b. Electrostatic interaction energy of UbcH5c with the H2A C-tail versus E2-NCP distance from all WT simulations. c. Electrostatic interaction energy of UbcH5c with the DNA versus E2-NCP distance from all WT simulations. The interaction energy during the initial 50 ns is highlighted in black. The initial energy of E2-H2A shows repulsive interactions that explains the effect of H2A C-tail on the upward motion of UbcH5c.

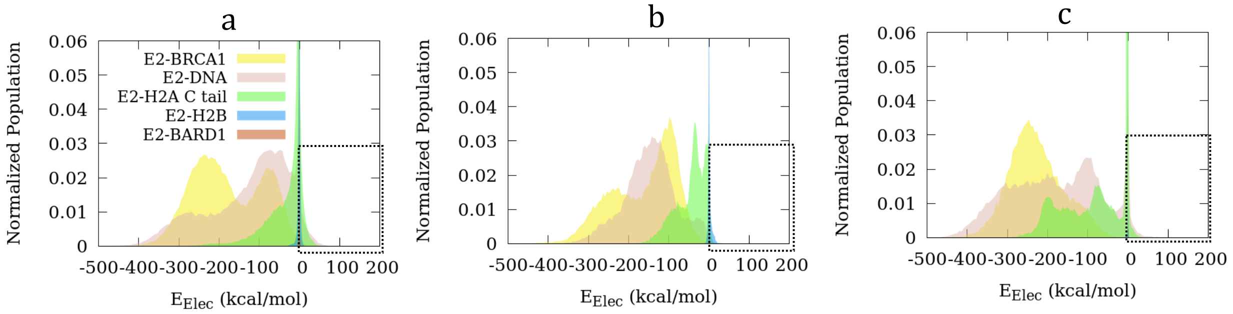


Figure S16: The electrostatic interaction energy of UbcH5c with H2A C-terminal tail, DNA, BRCA1, BARD1, and H2B for a. WT, b. all-ALA and c. all-ARG simulations.

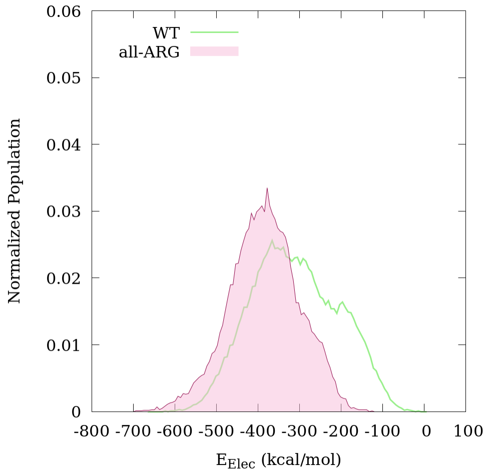


Figure S17: The electrostatic interaction energy between H2A C-tail and DNA in WT and all-ARG simulations.

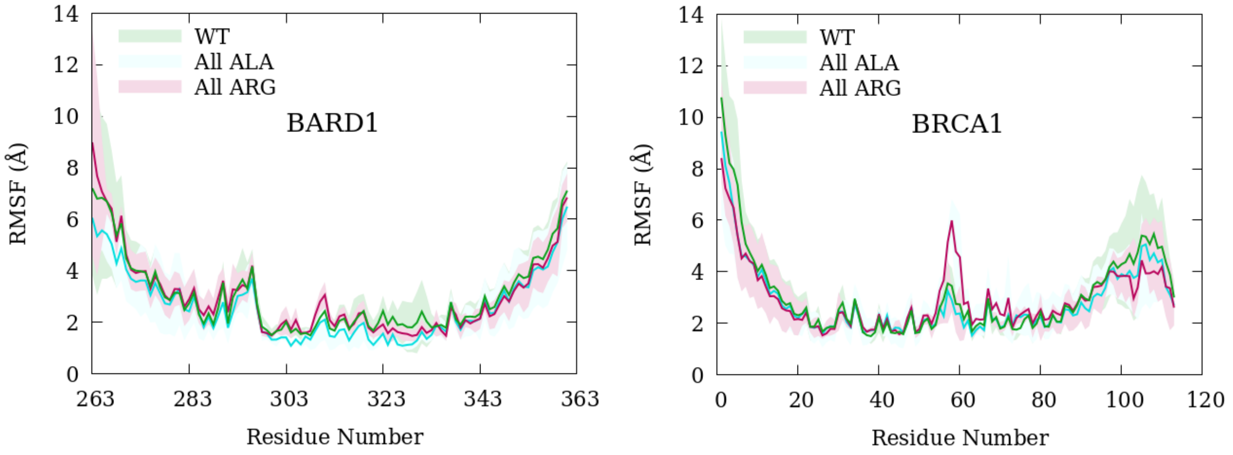


Figure S18: RMSF of BRCA1 and BARD1 obtained for WT, all-ALA and all-ARG mutant simulations.

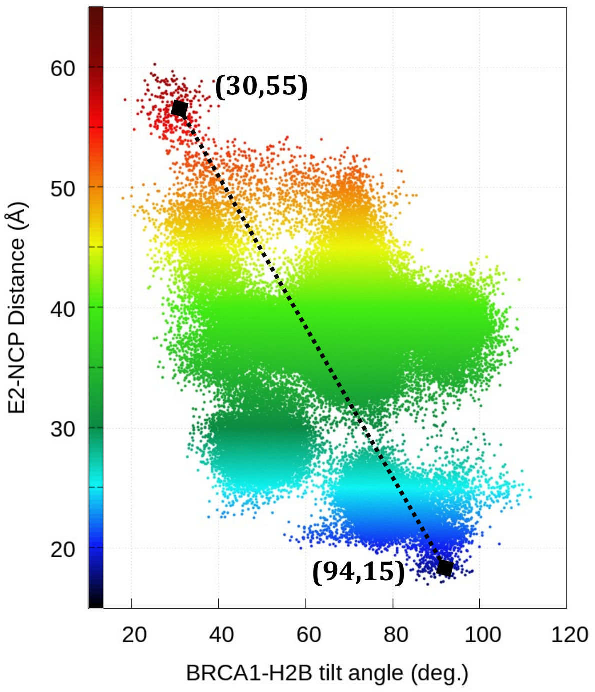


Figure S19: UbcH5c (E2) and NCP distance versus BRCA1-H2B tilt angle for WT simulations. Colored according to the values in Y axis (E2-NCP distance). Correlation of data shown in a dotted line.

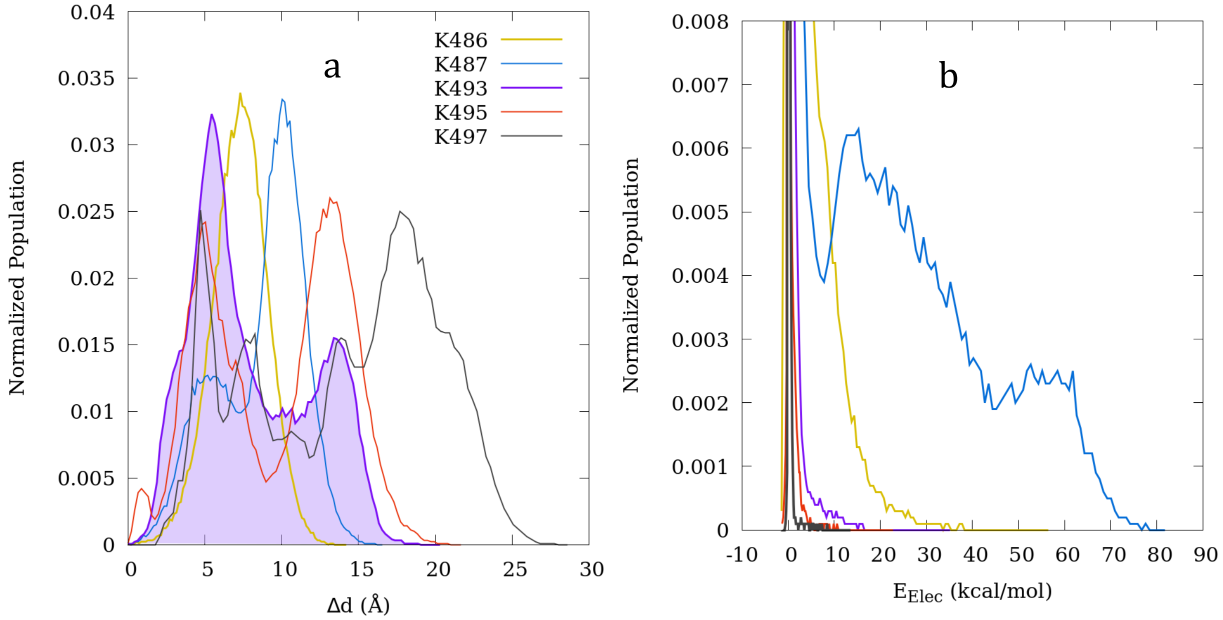


Figure S20: a. Normalized change in distance between 5 lysine residues ($C_\alpha$ atoms) in the H2A C-tail and P atoms of the 1443 DNA residue to explain the overall H2A C-tail motion. b. Normalized interaction energy between 5 lysine residues in the H2A C-tail and active site K197 for all WT simulations.

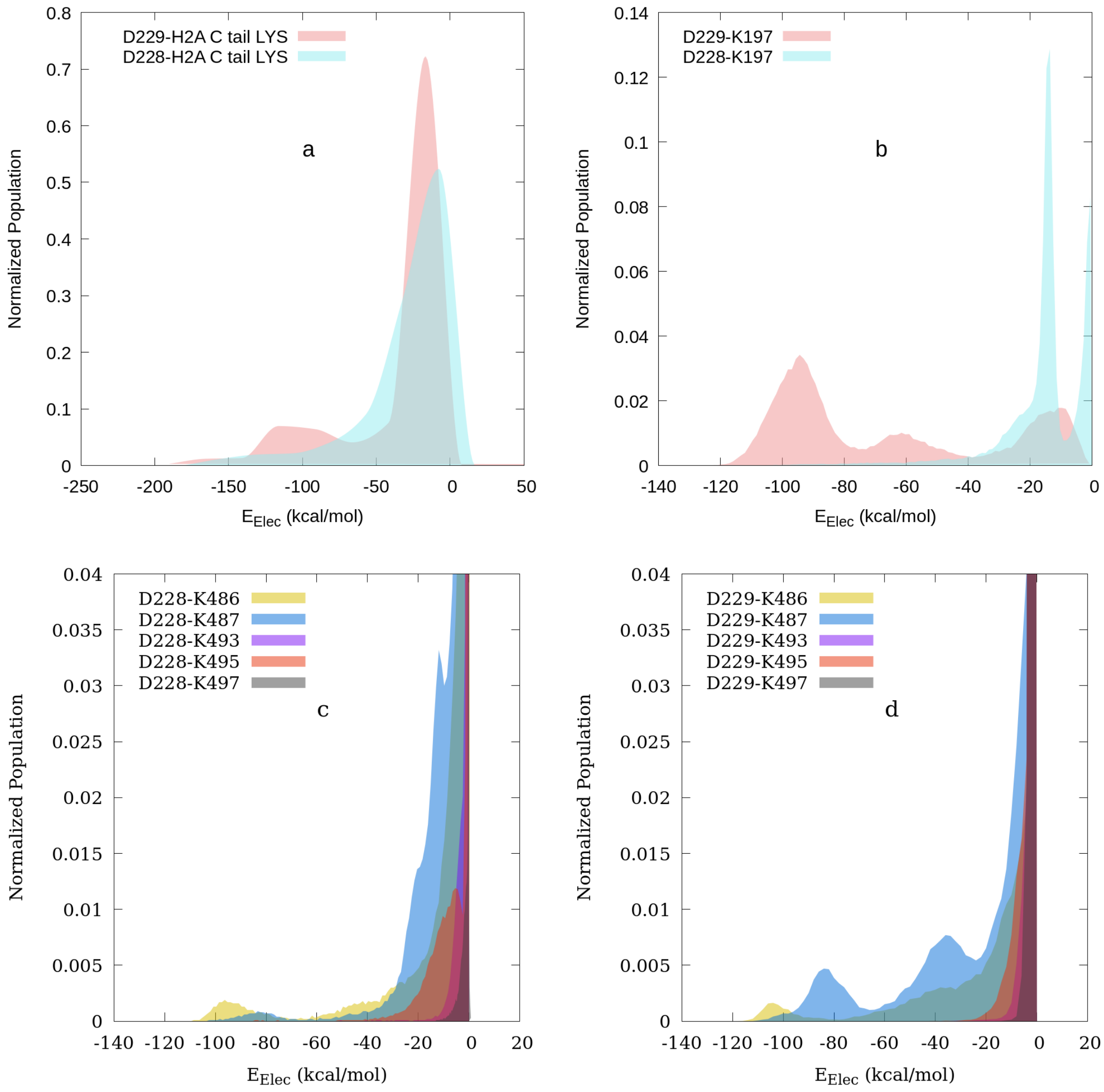


Figure S21: Electrostatic interaction energy between, a. UbcH5c D228 and D229 residues with H2A C-tail lysines. b. UbcH5c D228 and D229 residues with UbcH5c active site K197. c. D228 with H2A C-tail lysine residues individually. d. D229 with H2A C-tail lysines individually.
